# A fast *Myh* super enhancer dictates adult muscle fiber phenotype through competitive interactions with the fast *Myh* genes

**DOI:** 10.1101/2021.04.17.438406

**Authors:** Matthieu Dos Santos, Stéphanie Backer, Frédéric Auradé, Matthew Wong, Maud Wurmser, Rémi Pierre, Francina Langa, Marcio Do Cruzeiro, Alain Schmitt, Jean-Paul Concordet, Athanassia Sotiropoulos, Jeffrey Dilworth, Daan Noordermeer, Frédéric Relaix, Iori Sakakibara, Pascal Maire

## Abstract

The contractile properties of adult myofibers are shaped by their Myosin heavy chain (MYH) isoform content. We identify by snATAC-seq a 42kb super-enhancer (SE) at the locus regrouping the fast *Myh* (f*Myh*) genes. By 4C-seq we show that active f*Myh* promoters interact with the SE by DNA looping, leading to the activation of a single promoter per nucleus. A rainbow mouse transgenic model of the locus including the SE recapitulates the endogenous spatio-temporal expression of adult f*Myh* genes. *In situ* deletion of the SE by CRISPR/Cas9 editing demonstrates its major role in the control of associated f*Myh* genes, and deletion of two f*Myh* genes at the locus reveals an active competition of the promoters for the shared SE. Last, by disrupting the organization of f*Myh*, we uncover positional heterogeneity within limb skeletal muscles that may underlie selective muscle susceptibility to damage in certain myopathies.

## Introduction

Skeletal muscles constitute the most abundant organ in an adult human, about 40% of its total body mass. Most skeletal muscles are composed of a mixture of myofibers with distinct contractile, metabolic, resistance to fatigue properties, as well as differential vulnerability in pathophysiological situations ^1^. These different myofibers can be classified as slow or fast subtypes that selectively express genes responsible for their specific properties ^2–4^. The most widely used classification of myofibers types is based on their Myosin heavy chain (MYH) expression profile ^4–7^. MYH, one of the most abundant proteins present in adult myofibers, is a major determinant of myofiber speed of contraction. Each of the mammalian MYH isoform is coded by a specific gene and adult slow-type myofibers express *Myh7* (also known as *MyHCI*, *β* or *slow*), adult fast-type myofibers express *Myh2* (*MyHCIIA*), *Myh1* (*MyHCIIX*), *Myh4* (*MyHCIIB*) or *Myh13* (*MyHCeo*). During embryonic development *Myh7* and two specific fast *Myh* (f*Myh*) genes, *Myh3* (*MyHCemb)* and *Myh8 (MyHCperi)* are expressed ^8^.

The f*Myh (Myh3, Myh2, Myh1, Myh4, Myh8,* and *Myh13*) are organized as a cluster within a 350kb region on mouse chromosome 11 ^9^. The adult fast *Myh2, Myh1* and *Myh4* genes are expressed at a low level during embryogenesis and start to be expressed at a much higher level after birth ^8, 10–12^. The mechanisms controlling the robust coordinated expression of f*Myh* genes in the hundreds nuclei of a myofiber are not understood. Special regulatory elements called super enhancers (SE) have been shown to control high expression levels for cell lineage identity genes. These SE are composed of multiple enhancer elements spanning 10 to 50 kb of DNA and allowing efficient expression of associated genes ^13–17^. As identity genes expressed at high levels in specific fast myofiber subtypes, f*Myh* genes are good candidates to be controlled by a SE in the skeletal muscle lineage. The clustered organization and strict temporal regulation of the f*Myh* locus shows similarities with that of the human *β-globin* locus ^18^. At the *β*-*globin* locus a common regulatory sequence called locus control region (LCR) interacts dynamically with the different promoters within the locus to activate a single *Globin* isoform in erythroid cells ^19–21^. We hypothesized that a LCR/SE at the f*Myh* locus may coordinate the expression of selective f*Myh* genes in adult myofibers to finely control their identity.

To characterize the cis-regulatory elements required for the complex regulation of the specific f*Myh* genes we performed snATAC-seq and 4C-seq experiments with adult skeletal muscles and identified a 42kb opened chromatin region interacting in an exclusive manner with the activated f*Myh* promoter at the locus through 3D chromatin looping as revealed by 4C-seq experiments. A mouse rainbow transgenic line including this SE recapitulates the spatio-temporal expression of endogenous *Myh2*, *Myh1* and *Myh4* genes. We further show by CRISPR/Cas9 editing that *in situ* deletion of this 42kb SE region prevents expression of fetal *Myh8* and adult f*Myh* genes at the locus leading to fetal myofibers devoid of sarcomeres, unable to contract and precluding breathing at birth. We also tested the hypothesis of promoter competition for the shared SE, and show that absence of *Myh1* and *Myh4* leads to increased expression of *Myh2*, *Myh8* or *Myh13* in specific subregions of limb muscles. Altogether our studies suggest that the f*Myh* SE is responsible for the non-stochastic robust coordinated f*Myh* gene expression in the hundreds of body myonuclei present in adult myofibers. Analysis of the phenotype of all forelimbs and hindlimbs muscles in genetic perturbations within the f*Myh* locus reveals different categories of muscle susceptibility reminiscent of the selective muscle vulnerability observed in different neuromuscular diseases.

## Results

### Identification of a super enhancer acting as a locus control region in the f*Myh* locus

The majority of adult myofibers express a single *Myh* gene among the subfamily of fast *Myh4*, *Myh1*, *Myh2*, or slow *Myh7* genes. Fast muscles like the quadriceps are composed of myofibers expressing predominantly *Myh4* or *Myh1* genes while slow muscles like the soleus are composed of myofibers expressing predominantly *Myh7* or *Myh2* genes (Figure 1A-B). To identify the regulatory elements controlling the expression of f*Myh* genes, we performed snATAC-seq experiments with nuclei isolated from adult fast quadriceps and slow soleus ^10^. Myonuclei were classified based on the chromatin accessibility in the promoter and gene body of *Myh* genes (Figure 1C-D). In f*Myh* myonuclei (*Myh2, Myh1, and Myh4*), we observed 7 chromatin accessibility peaks in an intergenic region between *Myh3* and *Myh2.* This chromatin region is “closed” in nuclei that do not express f*Myh* genes like slow *Myh7* myonuclei and Fibro Adipogenic Progenitors (FAPs) nuclei where no snATAC-seq peak is detected (Figures 1D, S1A). These chromatin accessibility peaks cover the *Linc-Myh* gene ^22^, and end 25kb upstream of *Myh2* promoter (Figure S1B). Because of its large size of 42kb, this element could correspond to a conserved super enhancer (SE) controlling the f*Myh* genes of the locus in mammals (Figure S1B).

**Figure 1.**
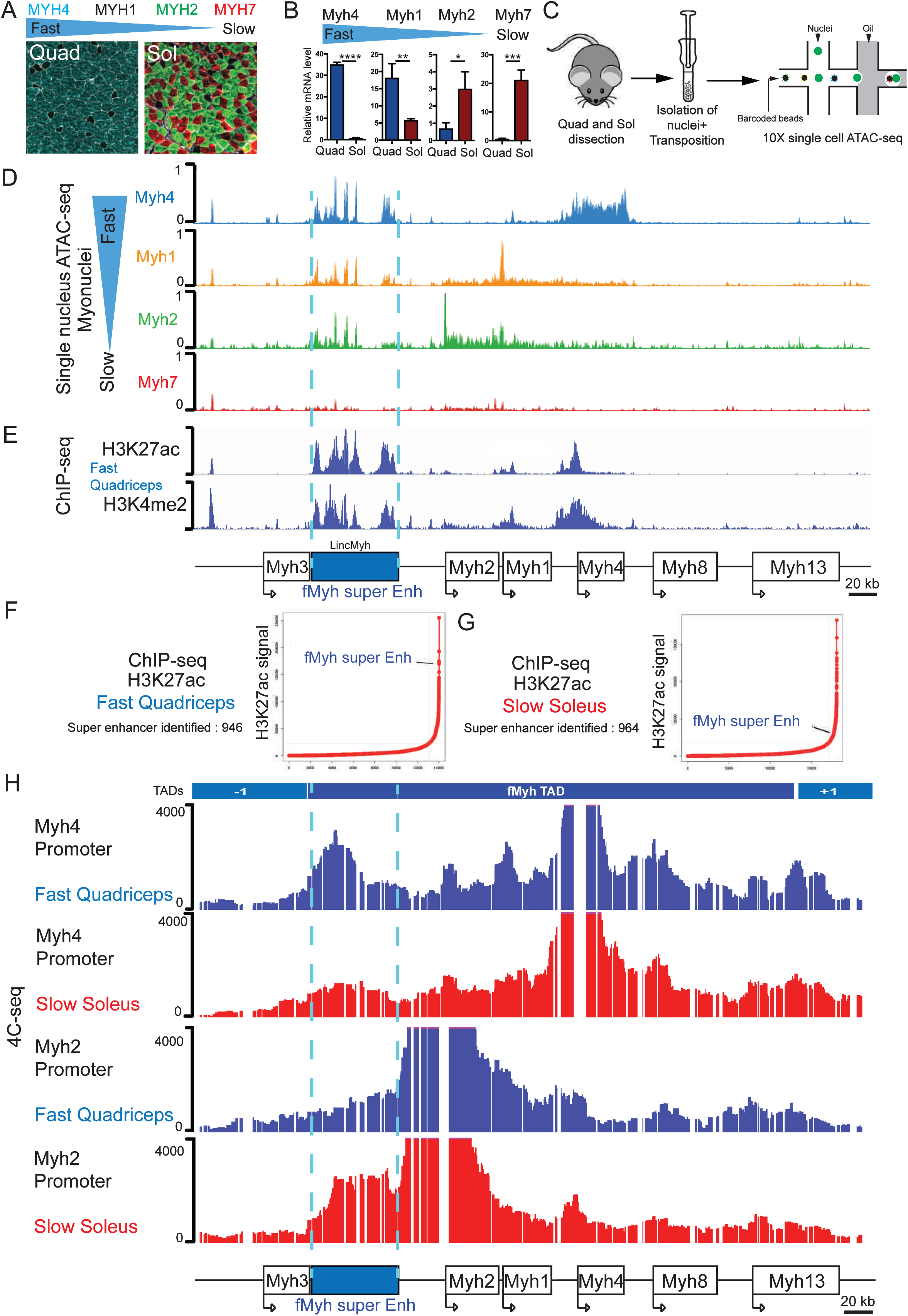
Identification of a super enhancer in the intergenic region of *Myh3* and *Myh2*. (A) Adult myofibers expressed different MYH isoforms. Immunostaining against fast (MYH4, MYH2) and slow (MYH7) MYH of adult fast quadriceps (Quad) and slow soleus (Sol) muscle sections, MYH1+ myofibers by default appear black. (B) Quantification by RT-qPCR of *Myh* mRNA expression in adult Quad and Sol. (C) Graphical scheme of the experiments used for snATAC-seq experiments performed with slow soleus and fast quadriceps adult skeletal muscle. (D) Chromatin accessibility of the different types of myonuclei in the fast *Myh* locus. In fast myonuclei, we identified a 42kb region with multiple chromatin accessibility peaks in the intergenic region of *Myh3* and *Myh2* genes. In slow *Myh7* myonuclei this region of chromatin is not accessible. (E) H3K27Ac and H3K4me2 ChIP-seq signals ^24^ were highly enriched in the 42kb region of snATAC-seq peaks in the intergenic region of *Myh3* and *Myh2* genes. (F) Plot showing ranked H3K27ac ChIP-seq signals for quadriceps enhancers. (G) Same as (F) in soleus. (H) 4C-seq experiments showing the interaction between the *Myh4* (Up) and *Myh2* (down) promoters in quadriceps and soleus: the promoter of the gene activated at the locus interacts with the 42kb cis-regulatory DNA module. Numerical data are presented as mean ± s.e.m. \**P* < 0.05, \*\**P* < 0.01, \*\*\**P* < 0.001, \*\*\*\**P* < 0.0001.

SE were first identified by Chip-seq by their higher level of transcription coactivators and active histone marks accumulation (H3K4me2 and H3K27ac) than conventional enhancers, and by their larger size compared with classic enhancers ^23^. They regulate the expression of highly transcribed genes specifying cell identity. To test if the element that we identified corresponds to a SE, we compared snATAC-seq results with available ChIP-seq against H3K4me2 and H3K27ac histone marks performed in skeletal muscle ^24^. We observed specific enrichment of these two active histone marks in the same snATAC-seq peaks of chromatin accessibility in the intergenic region between *Myh3* and *Myh2* in quadriceps and soleus (Figures 1E, S1A, ^24^). To determine if this sequence is a SE, we classified the slow and fast specific muscle active enhancers according to the enrichment in H3K27ac histone marks. The 42kb regulatory region of the *fMyh* locus shows a strong enrichment in H3K27ac marks compared to the other enhancers (Figure 1F). This showed that this 42kb intergenic region between *Myh3* and *Myh2* possesses the characteristics of a SE ^15^ that could control the expression of adjacent f*Myh* genes in the fast quadriceps but also in the slow soleus where around 50% of myofibers express *Myh2* or *Myh1* (Figures 1G, S1A). Based on these criteria, we called this regulatory element f*Myh*- SE.

One of the first SE discovered is the locus control region (LCR) of the *β-globin* locus ^25^. Like the f*Myh* locus, the human *β-globin* locus contains a cluster of *globin* gene isoforms expressed sequentially during embryonic, fetal, and adult erythropoiesis ^26^. The LCR of the *β-globin* locus forms dynamical and specific chromatin loops with the promoter of the gene transcribed at the locus. The similarities between clustered organization and temporal expression at the *β-globin* and f*Myh* loci suggested that the f*Myh*-SE could act by chromatin looping. To verify this, we performed 4C-seq by purifying nuclei from fast quadriceps and slow soleus. We designed specific primers to quantify the DNA regions interacting with *Myh4* and *Myh2* promoters when these genes are expressed or not. We observed stronger interactions between the *Myh4* promoter and the f*Myh*-SE in the quadriceps where this gene is more transcribed than in the soleus (Figure 1H). On the contrary, we observed more interactions between the *Myh2* promoter and the f*Myh*- SE in the soleus where this gene is more transcribed than in the quadriceps. We confirmed these results by quantifying the interactions between the f*Myh*-SE and other DNA regions in fast muscles. We observed strong and specific interactions between the f*Myh*-SE and the *Myh4* promoter in muscles expressing predominantly *Myh4* gene (Figure S2B). These results showed that the *fMyh*-SE forms specific and dynamic chromatin loops with the promoters of the genes transcribed at the locus, with 3D spatial proximity directly coinciding with activity of the promoter in the fiber type.

### The f*Myh* locus is organized in two topological associated domains (TADs)

In mammals, interactions between enhancers and promoters take place preferentially within TADs that are delimited by CTCF insulator binding sites. These CTCF sites can prevent enhancers from activating a gene present in another TAD ^27–30^. TAD organization and CTCF insulator sites are conserved between cells and mammalian genomes ^31^. We collected data of TAD organization in f*Myh* genes from available Hi-C experiments in embryonic stem cells ^32^. As shown in Figure S 2, the f*Myh* genes are clustered in two distinct TADs separated by CTCF binding on boundary elements observed in ChIP-seq experiments ^33^. One TAD includes the embryonically expressed *Myh3*, and another adjacent TAD includes all the other *fMyh* genes. To confirm this 3D organization of the fast *Myh* locus, we performed 4C-seq experiments with different viewpoints all along the locus. These experiments confirmed that the *Myh3* promoter interacted mostly with DNA sequences present in its TAD (TAD-1), while the other f*Myh* promoters interacted almost exclusively with sequences present in the f*Myh* TAD. We also observed that the f*Myh*-SE interacted mostly with sequences present in the f*Myh* TAD (Figure S2B). This suggesting that the adult f*Myh* genes, the fetal *Myh8* gene and the extraocular muscle-specific *Myh13* gene, that are all located in the same TAD, could be controlled by the same f*Myh*-SE. On the contrary, either the regulatory element(s) that control the spatio- temporal expression of *Myh3* should be distinct from the ones controlling the other f*Myh* genes or the TAD boundary should be dynamically reorganized in cells where this gene is active.

### A transgenic mouse model of the f*Myh* locus fully recapitulates *Myh1*, *Myh2,* and *Myh4* expression

To create a transgenic mouse model for f*Myh* expression, we inserted into a 222kb bacterial artificial chromosome (BAC) that partially covered the f*Myh* locus (end of *Myh3* to the middle of *Myh8*), the cDNAs coding for YFP at the ATG of *Myh2*, Tomato at the ATG of *Myh1,* and CFP at the ATG of *Myh4* to test their correct expression. A stop codon and a polyA tail were also inserted at the end of each transgene, preventing the expression of fusion proteins between cDNAs and the associated f*Myh*. The recombined BAC was injected in mouse oocytes and 2 separate transgenic animals were obtained and analyzed. We determined by qPCR on genomic DNA that one transgenic line called Enh+ integrated 2 complete copies of the entire length of the BAC including the SE. The second independent mouse line called Enh-, possesses an incomplete copy of the BAC devoid of the f*Myh* SE (Figure 2A). We observed efficient YFP, Tomato, and CFP expression in all skeletal muscles of Enh+ animals (Figures 2B-D, S3A). Expression of the transgenes was not detected in the lung, liver, heart, or kidney (Figure S3A). Next, we compared the expression of the three transgenes with the accumulation of endogenous MYH proteins and mRNAs. As seen in Figure 2E, YFP myofibers were detected in the slow soleus, in agreement with endogenous MYH2 expression, Tomato myofibers were detected in bracoradial muscles, and CFP myofibers in the quadriceps. By immunohistochemistry we observed a strong correlation between the expression of endogenous MYH2 proteins and YFP+ myofibers, and between MYH1 proteins and Tomato myofibers (Figure 2F-G). We did not observe the expression of the transgenes in slow MYH7 myofibers of the soleus (Figure S3B). This correlation between transgene and endogenous gene expression was confirmed by RT-qPCR. Efficient *YFP* and *Myh2* mRNA accumulation was found in the soleus of Enh+ mice. *Tomato* mRNAs accumulated in both quadriceps and soleus like *Myh1* mRNA. *CFP* mRNA accumulated more in quadriceps than in soleus like *Myh4* mRNAs (Figure 2F-H). The three transgenes were detected in all skeletal muscles of the body including extraocular muscles and Esophagus (Figure S4A,B). Tomato expression was first detected at P0 in the diaphragm when corresponding f*Myh* genes expression become detected (Figure S4C) ^10^. Interestingly, most adult Enh+ myofibers expressed only one transgene although hybrid fibers ^34^ were also observed (Figure S4D-F).

**Figure 2.**
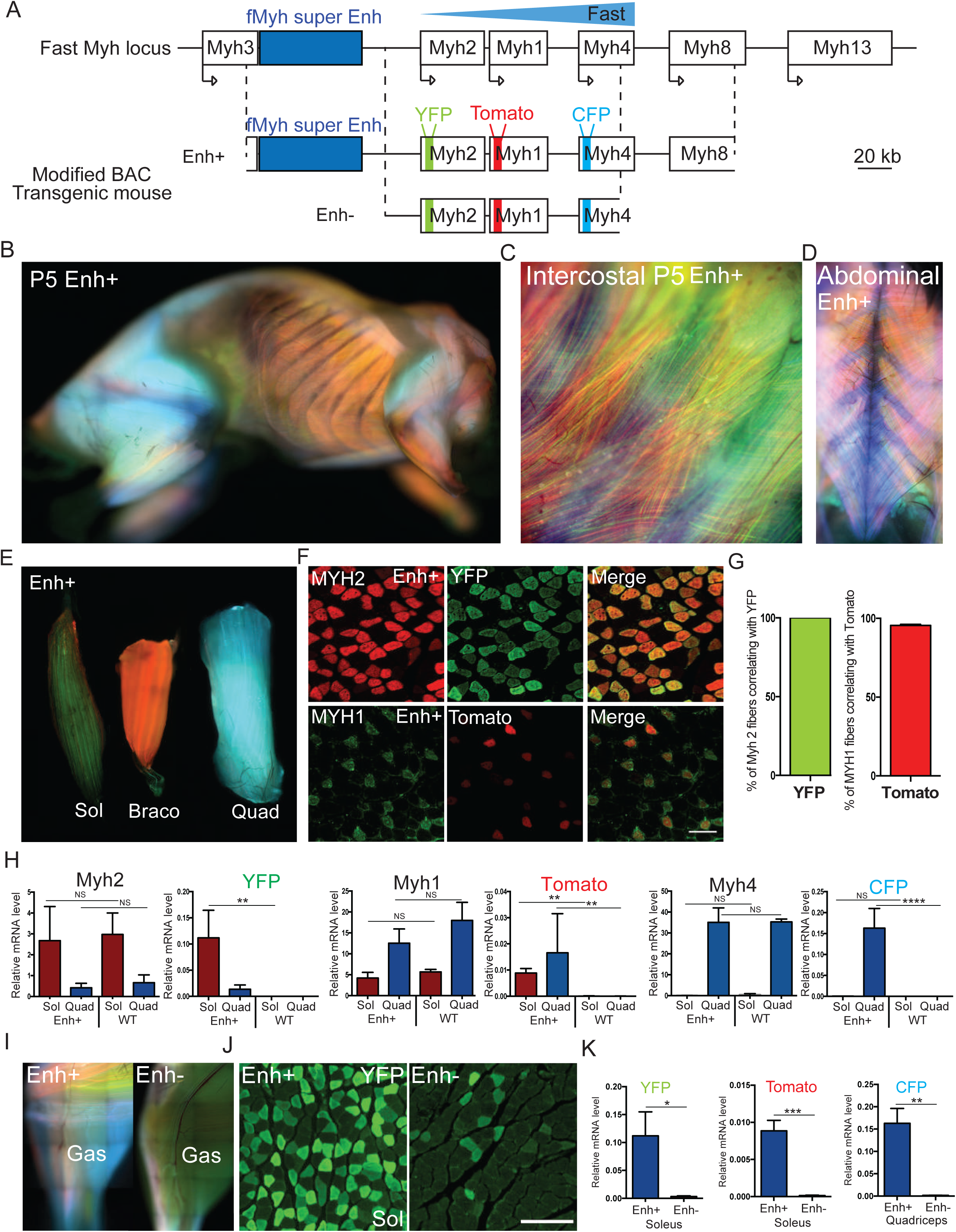
Transgenic models to study f*Myh* genes expression. (A) Schematic representation of mouse f*Myh* locus and the recombined 222kb Bacterial Artificial Chromosome (BAC) of the same locus. YFP, Tomato and CFP cDNAs were inserted in the first exon of *Myh1*, *Myh2* and *Myh4* genes respectively in the BAC. Two transgenic mouse lines were obtained, one called Enh+ that integrated 2 complete copies of the BAC and the other called Enh- devoid of the SE region and the 3’ region of the locus. The transgenes YFP, Tomato and CFP are not to scale. (B) Picture of a five days old Enh+ transgenic mouse, lateral view. Red; Tomato, green; YFP and blue; CFP. All skeletal muscles expressed the transgenes. (C) Same as (B), zoom in intercostal muscles. (D) Same as (B) in intercostal and abdominal muscles of a 2-month-old Enh+ transgenic mouse. (E) Transgene expression in adult soleus (Sol), bracoradial (Braco) and quadriceps (Quad) showing predominant expression of YFP in green, Tomato in red and CFP in blue for each muscle. (F) Expression of the transgenes correlates with endogenous MYH protein expression in Enh+ line. Up: immunofluorescence against endogenous MYH2 (red) and of YFP (green) in adult soleus transverse section of Enh+ mice. Down: immunofluorescence against endogenous MYH1 (green) and of Tomato (red) in adult quadriceps transverse section of Enh+ mice. (G) Quantification of the percentage of MYH2 or MYH1 fibers expressing YFP or Tomato respectively, (n=3). All MYH2 fibers are YFP+ and almost all MYH1 fibers are Tomato+. (H) Relative expression level of mRNA in adult Sol and Quad of endogenous *Myh* genes and of transgenes, in wild type (wt) and in Enh+ mice (n=3). (I) Pictures of the adult leg of Enh+ (left) and Enh- (right) mouse. The expression of the three transgenes is much higher in the Enh+ line compared to Enh- mouse. (J) Immunostaining with GFP antibodies revealing YFP fibers on a section of adult Sol in Enh+ and Enh- mice. In Enh+ mouse, all MYH2 fibers expressed YFP whereas in Enh- only 10% of MYH2 fibers expressed YFP. (K) Quantification by RT-qPCR of transgenes expression in Enh+ and Enh- mouse line. Numerical data are presented as mean ± S.E.M. \**P* < 0.05, \*\**P* < 0.01, \*\*\**P* < 0.001. Scale bars: 100μm for F, and 50μm for J.

While efficient expression of CFP, Tomato and YFP was detected in the skeletal muscles of Enh+ animals, very low expression of the three transgenes was observed in Enh- animals (Figure 2I). This decreased transgene expression was also observed by immunostaining on adult muscle sections: much fewer YFP fibers in soleus and much fewer CFP fibers in gastrocnemius were detected in Enh- mice as compared to Enh+ mice (Figures 2J, S5A). Transgenes mRNA level was at least 100- fold lower in Enh- than in Enh+ animals, as estimated by RT-qPCR (Figure 2K). Altogether, our results show that all regulatory sequences to fully recapitulate the spatiotemporal expression patterns of the f*Myh* genes are present in the modified 222kb BAC in Enh+ mice, which roughly overlaps the f*Myh* TAD, and that the f*Myh*-SE and/or other sequences absent in Enh- transgenic animals are required for efficient *Myh2-YFP*, *Myh1-Tomato* and *Myh4-CFP* transgenes expression. Lastly the Enh+ rainbow mouse line allows visualizing the fiber-type switches occurring during denervation, aging, in muscle specific *Six1* conditional knock out mouse models, and in other conditions at an individual scale (Figure S6A-F) and is thus a powerful tool to study fiber-type changes in pathophysiological conditions ^4, 5^ .

### The f*Myh*-SE is required for adult f*Myh* and neonatal *Myh8* expression

To assess the requirement of the SE for efficient f*Myh* genes expression *in vivo,* we generated by CRISPR/Cas9 genome editing a knock-out mouse line deleted of this 42kb element (Figures 3A, S7A-B). Heterozygote mutant mice were viable and fertile and presented no obvious deleterious phenotype. In contrast, homozygote mutants died at birth, potentially due to impairment of respiratory skeletal muscle contractions as suggested by the absence of air in their lungs (Figure 3B). E18.5 mutant fetuses showed no major visible skeletal muscle hypoplasia (Figures 3C, G, S7C). In muscles of E18.5 mutant fetuses, the deletion of the f*Myh*-SE induced a strong decrease of the expression of adult f*Myh* (*Myh2, Myh1,* and *Myh4*) and of neonatal *Myh8* genes detected by RNAscope on isolated fibers from the diaphragm and quantified by RT-qPCR on leg skeletal muscles (Figure 3D-F). We detected at this embryonic stage regionalized low expression of adult f*Myh* along a few mutant fibers (Figure S 7D), indicating that the *Myh4* gene can be expressed in rare myonuclei in absence of the f*Myh-*SE. Thus, the f*Myh-*SE allows sustained expression of f*Myh* in the syncytium, although not all myonuclei at E18.5 have yet activated the expression of these adult forms (Dos Santos, 2020). *Myh4* or *Myh1* simple KO ^35, 36^ and *Myh1*/*Myh4* double KO (see below) mouse are viable and fertile. *Myh2* mutant mice have not been analyzed in detail but seem to have no major phenotype ^37^. The absence of breathing and survival observed in P0 42kb f*Myh*-SE mutants could be due to the loss of *Myh8* expression, or to the loss of the expression of a combination of several f*Myh* genes (Figure 3D, G, H). At the limb level expression of *Myh3* and *Myh7* was not affected as shown by RT-qPCR experiments and by immunocytochemistry against MYH3 and MYH7 (Figures 3G, H, S7C). In the 42kb f*Myh*-SE E18.5 mutants many limb myofibers presented absence of sarcomeres associated with Actin aggregates around their myonuclei with only a few fibers that did not present these defects (Figures 3J, S7E). Electronic microscopy experiments showed an accumulation of fibrillar materials in mutant diaphragm myofibers that may correspond to Actin accumulation in absence of MYH proteins (Figure 3K), and the absence of sarcomere in many myofibers. These defects of sarcomere formation in mutant myofibers did not impair their innervation but seemed to affect neuromuscular junctions distribution in the diaphragm (Figure 3I). We suspect that unaffected fibers could be primary fibers expressing *Myh7* and or *Myh3*, whose expression appeared normal, while affected myofibers could be secondary myofibers that normally activate the expression of *Myh8* (Figure 3E). The absence of MYH8 could thus lead to sarcomere formation defects leading to Actin aggregates. Altogether these results showed that the f*Myh* SE controls the expression of adult f*Myh* and neonatal *Myh8* and that these isoforms are required for correct sarcomere formation in secondary myofibers and important for efficient muscle contraction at birth.

**Figure 3.**
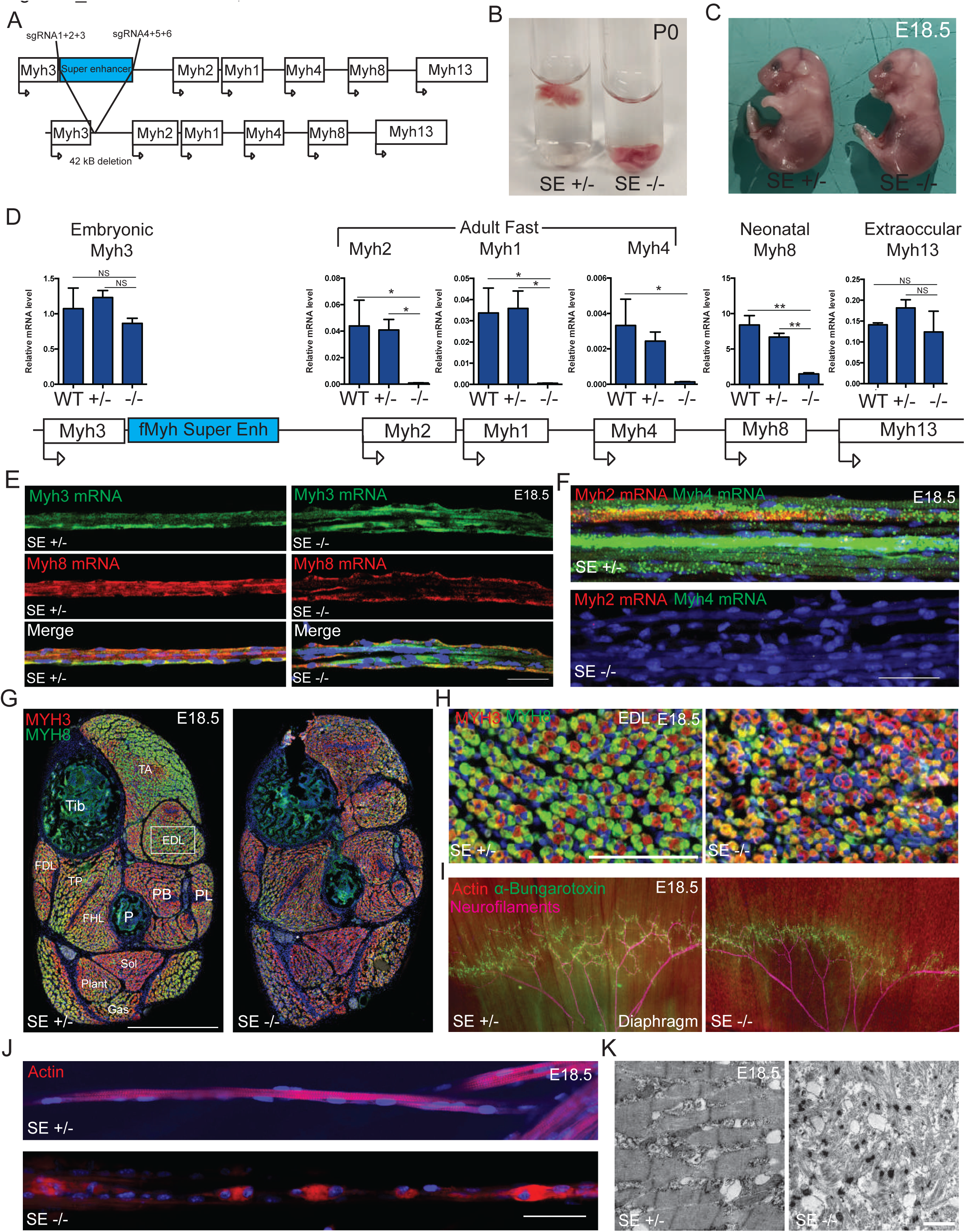
The f*Myh*-SE is required for adult f*Myh* and neonatal *Myh8* genes expression. (A) A mouse line deleted for the f*Myh*-SE element was generated by injecting specific sgRNAs and Cas9 protein into mouse oocytes. (B) f*Myh-*SE^-/-^ mice died at birth (P0) without breathing and air in their lungs. (C) f*Myh-*SE^-/-^ E18.5 fetuses did not present severe visible malformations. (D) Quantification of *Myh* mRNAs by RT-qPCR in control and f*Myh-*SE^-/-^ E18.5 forelimb skeletal muscles. Mutant muscles showed decreased *Myh2*, *Myh1*, *Myh4* and *Myh8* mRNAs levels. (E) RNascope experiments against *Myh3* and *Myh8* mRNAs on isolated E18.5 forelimb fibers of control and mutant mice. (F) Same as (E) showing a decreased accumulation of *Myh2* and *Myh4* mRNAs in mutant mice compared to their littermate controls. (G) Immunostaining at the distal hindlimb level of E18.5 control and mutant fetuses revealing MYH3 and MYH8 positive myofibers. (H) Same as (G), zoom in the EDL of control and mutant. (I) *In toto* immunostaining of diaphragms from E18.5 mutant and control fetuses showing in red Actin filaments (phalloidine), in green AchR (alpha-bungarotoxin), and in pink neurofilaments showing altered repartition of NMJ and punctated Actin aggregates in mutant diaphragms. (J) Myofibers from mutant diaphragm showed defects in sarcomeres organization as shown by phalloidine staining. (K) Electronic microscopy pictures of the sarcomeres defects present in mutant E18.5 fetuses compared to their littermate controls. For D (n=3). For E and F, scale bar: 50 μm. For G, scale bar: 500 μm. For H, scale bar: 100 μm. Numerical data are presented as mean ± s.e.m. \**P* < 0.05, \*\**P* < 0.01.

### The f*Myh*-SE is composed of distinct cis-regulatory modules (CRM)

SEs are composed of multiple enhancers and each with a specific role in promoter activation ^38, 39^. To characterize the role of two individual CRM identified by snATAC-seq experiments in the SE, we generated their deletion by CRISPR/Cas9 genome editing and obtained two distinct mouse mutant lines. The first CRM enhancer A (*EnhA*) corresponds to a 5Kb region located at the most 3’ snATAC-seq peaks of the f*Myh*-SE (Figure 4A). The second CRM enhancer B (*EnhB*) corresponds to two snATAC-seq peaks located in the middle of the f*Myh*-SE. We previously showed that this CRM can activate the expression of *Myh1*, *Myh2,* and *Myh4* promoters in transient adult muscle transfection assays ^22^. In contrast to homozygote mice deleted for the f*Myh*-SE that died at birth, we obtained viable and fertile adult *EnhA* and *EnhB* homozygote mutant mice. We determined the expression of MYH7, MYH2, and MYH4 in the distal hindlimb by immunohistochemistry (Figure 4B) of these mutants. *EnhA^-/-^* mice showed a strong decrease of MYH4 expression in certain specific muscles (Figure 4B-C). MYH4 was no more detected in the TP and the FHL limb muscles of *EnhA^-/-^*, while the number of MYH1 fibers increased in these mutant muscles (Figures 4B-D, S8D). This MYH4 fiber-type switch associated with the absence of the *EnhA* was also observed in other muscles (TA, EDL, PB, PL, FDL, and Plant) while other muscles (Gas and Sol) were spared. This result was confirmed by RT-qPCR experiments showing downregulation of *Myh4* expression in the TA of *EnhA* mutants (Figures 4E, S8E). These results showed that enhancer A dominates regulation of *Myh4* in specific muscles, probably through the recruitment of key *Myh4* identity factors, while dispensable in others and showed also that MYH4 myofibers are not all equivalent. A low expression of *Myh8* and *Myh13* was also detected in adult WT TA which was strongly decreased in *EnhA^-/-^* TA, demonstrating that the expression of these two genes is also controlled by the enhancer A present in the SE (Figure 4E).

**Figure 4.**
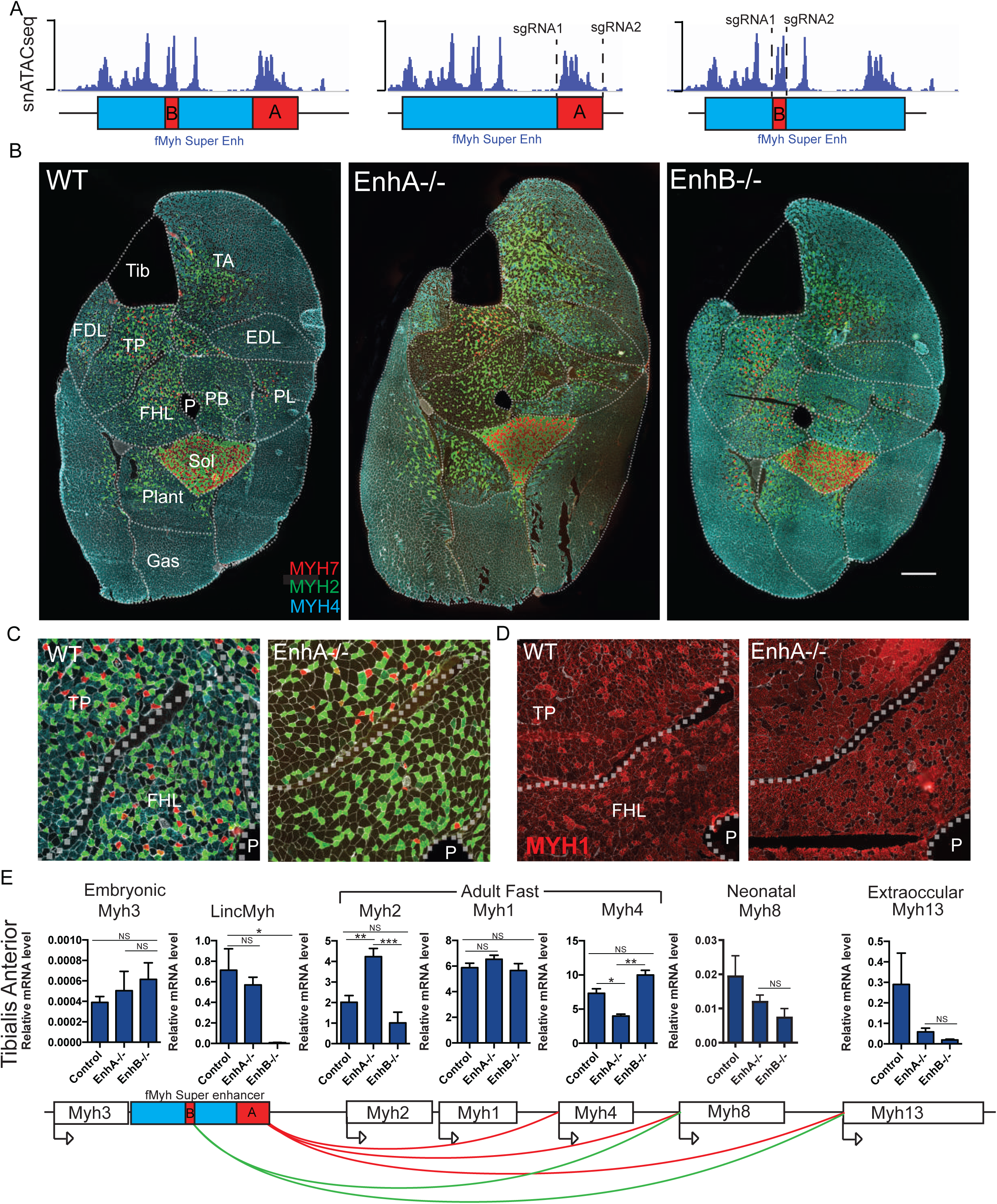
Role of the different enhancers composing the SE. (A) Schematic representation of the snATAC-seq peaks along the 42kb SE and the enhancers A and B deleted by CRISPR/Cas9 editing. (B) Immunostaining against fast MYH2, MYH4 and slow MYH7 on adult leg sections of 2-3-month-old mouse female deleted for enhancer A or B. (C) Same as (B), zoom in Tibialis posterior and FHL muscle of WT and *EnhA^-/-^* mutant. (D) Immunostaining against fast MYH1 in Tibialis posterior and FHL muscle of WT and *EnhA^-/-^* mutant. The absence of *EnhA* induced an increased number of MYH1 positive fibers. (E) Quantification of f*Myh* mRNA and of *Linc-Myh* in adult TA of control and *EnhA* and *EnhB* mutant by RT-qPCR experiments. Schema of the interactions identified. For E, n=3. Numerical data are presented as mean ± S.E.M. \**P* < 0.05, \*\**P* < 0.01, \*\*\**P* < 0.001. Scale bars: 100μm for B.

In contrast to the *EnhA* mutant mice, we observed no major modification of slow MYH7 and fast MYH2 and MYH4 expression in muscles of *EnhB* mutant animals by immunostaining (Figure 4B). As for *EnhA* mutant muscles, we observed a decrease of *Myh8* and *Myh13* expression in adult *EnhB* mutant TA compared to the WT (Figure 4E). *Linc-Myh* expression was no more detected in *EnhB^-/-^* TA. We further generated a transgenic mouse line carrying an *nls-LacZ* transgene under the control of *EnhB* DNA sequences (Figure S8F-G). *Nls-LacZ* transgene expression was detected only in fast and not in slow muscles. These results showed that even if the deletion of *EnhB* do not induce major alterations of adult f*Myh* expression, this DNA element has an enhancer activity in fast adult fibers, as already suggested ^22^. Altogether analysis of these mutant mouse lines revealed that the SE is composed of distinct enhancer elements possessing distinct functions, two of which activate *Myh1*, *Myh2*, *Myh4*, *Myh8 or Myh13* genes in specific muscles.

### The f*Myh* gene promoters compete for the SE

To further elucidate the mechanisms controlling the specific and exclusive activation of f*Myh* promoters, we tested whether these promoters competed for the SE. A mouse model harboring a 72kb deletion of the *Myh1* and *Myh4* genes (*Myh(1-4)^Del^*) was generated by CRISPR/Cas9 genome editing (Figures 5A, S9A-B). In the deleted allele, *Myh8* and *Myh13* genes are brought closer to the f*Myh-*SE, while the *Myh2* promoter remains at the same distance from the f*Myh-* SE than in the wt allele. M*yh(1-4)^Del/+^* and *Myh(1-4)^Del/Del^* animals were viable. No expression of *Myh1* and *Myh4* was detected in *Myh(1-4)^Del/Del^* animals. These mutants presented a strong hypotrophy in specific areas of individual skeletal muscles, while other areas of the same muscle seemed preserved: the deeper regions of the TA and Gas were more spared than the superficial regions where small myofibers accumulated (Figure 5B). This selective partitioning seemed to less affect deep muscles (Plantaris, PB) compared to the superficial areas of peripheral muscles like the TA or the Gas (Figure 5B). In *Myh(1-4)^Del/+^* and *Myh(1-4)^Del/Del^* mouse, we observed increased *Myh2* expression showing that the deleted allele for *Myh1* and *Myh4* does ectopically activate *Myh2* in the deep regions of muscle masses (Figures 5B, E, S9E-F). We also detected an increased expression of *Myh8* and *Myh13* in both *Myh(1-4)^Del/+^* and *Myh(1-4)^Del/Del^* mutant muscles (Figure 5C, D, E). Interestingly we observed in this *Myh(1- 4)^Del/Del^* mutant a deep to peripheral gradient of MYH8 and MYH13 positive myofibers, with increased MYH13 fibers at the peripheral regions of muscle masses (Figure 5C, D). Thus, in absence of *Myh1* and *Myh4* genes, the f*Myh*-SE can activate the expression of either *Myh2*, *Myh8* or *Myh13,* with a degree of plasticity of the myofibers depending on their position inside each individual muscle. These results show that each f*Myh* promoter competes for interaction with the SE and that this competition is influenced by specific muscle subvolumes in agreement with a selective partitioning ^40^, and by the deep or superficial position of the muscle itself.

**Figure 5.**
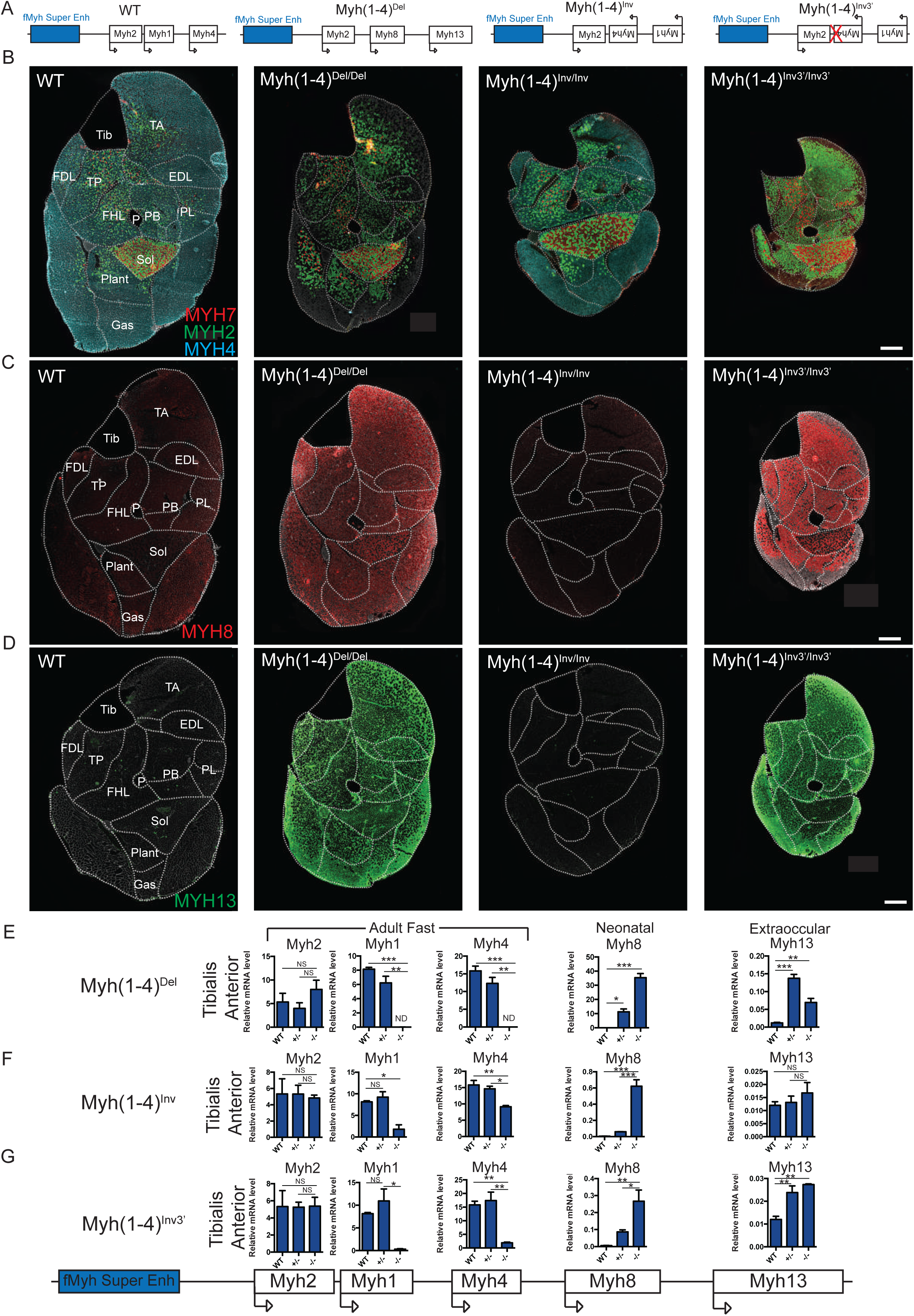
The promoters of f*Myh* genes compete for the shared SE. (A) Schema of the distinct f*Myh* alleles generated by CRISPR/Cas9 editing. (B) Immunostaining against MYH4 (blue), MYH2 (green) and slow MYH7 (red) of adult distal hindlimb sections of wt, *Myh(1-4)^Del/Del^*, *Myh(1-4)^Inv/Inv^* and of *Myh(1-4)^Inv3’/Inv3’^* mutants. (C) Immunostaining against neonatal MYH8 of adult leg sections in wt, *Myh(1-4)^Del/Del^*, *Myh(1- 4)^Inv/Inv^* and of *Myh(1-4)^Inv3’/Inv3’^*. (D) Same as (C) against extraocular MYH13. (E) Quantification of *Myh2, Myh1, Myh4, Myh8 and Myh13* mRNAs of adult wt, *Myh(1-4)^Del/+^* and *Myh(1-4)^Del/Del^* TA by RT-qPCR experiments. (F) Quantification of *Myh2, Myh1, Myh4, Myh8 and Myh13* mRNAs of adult wt, *Myh(1-4)^Inv/+^* and *Myh(1-4)^Inv/Inv^* TA by RT-qPCR experiments. (G) Quantification of *Myh2, Myh1, Myh4, Myh8 and Myh13* mRNAs of adult wt, *Myh(1-4)^Inv3’/+^* and *Myh(1-4)^Inv3’/Inv3’^* TA by RT-qPCR. For E, F, G (n=3). Numerical data are presented as mean ± S.E.M. \**P* < 0.05, \*\**P* < 0.01, \*\*\**P* < 0.001. Scale bars: 50μm for G.

With the sgRNA used to delete *Myh1* and *Myh4,* we obtained two additional mouse lines with a complete inversion of *Myh1* and *Myh4* genes (*Myh(1-4)^Inv^* and *Myh(1-4)^Inv3’^*), allowing to test the hypothesis that the order of the *Myh1* and *Myh4* genes in the locus is important for their correct expression. In both these lines, the order of the f*Myh* genes in the locus was modified (*Myh2*, *Myh4*, *Myh1* then *Myh8*). The homozygote *Myh(1-4)^Inv/Inv^* mutant mice were viable at the homozygous state and showed a strong decrease of *Myh1* expression, a weaker decrease of *Myh4* expression and no difference of *Myh2* expression compared to WT mice (Figure 5B-F). This indicates that a closer proximity of the *Myh4* promoter to the SE did not increase its activity at the adult stage. The strong decrease of *Myh1* expression could be due to the increased distance between its promoter and the f*Myh*-SE, to the misorder of the genes at the locus, or more probably to missing elements in the *Myh1* promoter, since only 575bp upstream of the transcription start site is associated with *Myh1* promoter in the inverted allele. We also observed an upregulation of *Myh8* in this mutant line (Figure 5F).

In the other *Myh(1-4)^Inv3’^* line, a deletion at the 3’ end of *Myh4* was observed, precluding MYH4 synthesis. The homozygote *Myh(1-4)^Inv3’/Inv3’^* mutant mice were viable, but presented a severe skeletal muscle atrophy. In this mutant mouse line, we observed a strong decrease of *Myh1* and *Myh4* expression (Figure 5B, G). Quantification of *Myh1* and *Myh4* pre-mRNA levels showed that the transcription at the *Myh4* gene in TA was modestly decreased in *Myh(1-4)^Inv/Inv^* and in *Myh(1-4)^Inv3’/Inv3’^* mutant as compared with WT, while *Myh1* transcription level was severely downregulated (Figure S9I-J). This showing that *Myh4* promoter can act as a decoy for the SE in *Myh(1-4)^Inv3’/Inv3’^*since no MYH4 protein is produced. Similarly to the *Myh(1-4)^Del/Del^* mouse line, we observed an upregulation of *Myh8* and *Myh13* expression in *Myh(1-4)^Inv3’/Inv3’^* muscles (Figure 5G). Interestingly in *Myh(1-4)^Inv3’/Inv3’^* animals we observed many MYH2/MYH8 hybrid fibers and many pure MYH13 fibers preferentially in superficial areas of peripheral muscles like the TA or the Gas. MYH13 positive fibers were atrophic (Figures 5D, 6C). We failed to detect MYH3 on *Myh(1-4)^Inv3’/Inv3’^* adult hindlimb sections (not shown). These results showed that in the inverted allele, the SE could activate misoriented *Myh4* gene, but less efficiently, and activated the expression of *Myh8* and *Myh13* in the myofibers. Altogether these results suggested that competition between the different *Myh* promoters for a shared SE controls their activation and that the order of the genes at the locus does not dictate their correct spatial expression.

### Limb skeletal muscles can be classified into 3 major categories with specific genetic programs

At least 640 different skeletal muscles can be identified in the human body, each with a specific form, architecture, position, and function. In several myopathies, skeletal muscles can be specifically affected depending on their anatomic position ^1^. Distinct genetic programs controlling the identity of each skeletal muscle in its specific environment may determine this selective vulnerability. The different mutants that were generated in this study presented distinct muscle phenotype depending on their location in the body. By comparing the fiber-type composition and fiber size in WT, *EnhA^-/-^* and *Myh(1-4)^Inv3’/Inv3’^* mutant mice (Figure 6A-C), we identified 3 different categories of skeletal muscles. The first category corresponded to muscles like the soleus, principally composed of small MYH7 and of MYH2 fibers (Figure 6A). The soleus muscle was not affected in *EnhA^-/-^* and *Myh(1-4)^Inv3’/Inv3’^* mouse. The second category of muscles included muscles similar to the Tibialis posterior principally composed of MYH2, MYH1 and MYH4 fibers (Figure 6B). These muscles were affected in *EnhA^-/-^* and *Myh(1- 4)^Inv3’/Inv3’^* mutant mice and did not express MYH4 anymore. The last category of muscle regrouped muscles similar to the gastrocnemius expressing mainly MYH4 (Figure 6C). The fibers of these groups of muscles presented a drastic decrease of fiber cross section area in the *Myh(1-4)^Inv3’/Inv3’^* mutants. In contrast, these muscles were not affected in *EnhA^-/-^* mice. We next extended this study in proximal and distal muscles of the fore- and hindlimbs (Figure 6D-G). As observed at the distal hindlimb level, muscles in forelimbs and proximal hindlimb showed distinct phenotype depending on their deep or superficial position^41^. We could detect specific localization of these 3 groups of muscles in the different parts of the hindlimb and forelimbs but with spatial patterns that seemed similar. The category of muscles with similar properties to the Soleus (shown in red) was the most internal in the limb. In contrast, the category of muscles with similar properties to the Gastrocnemius (shown in blue) was the most external.

**Figure 6.**
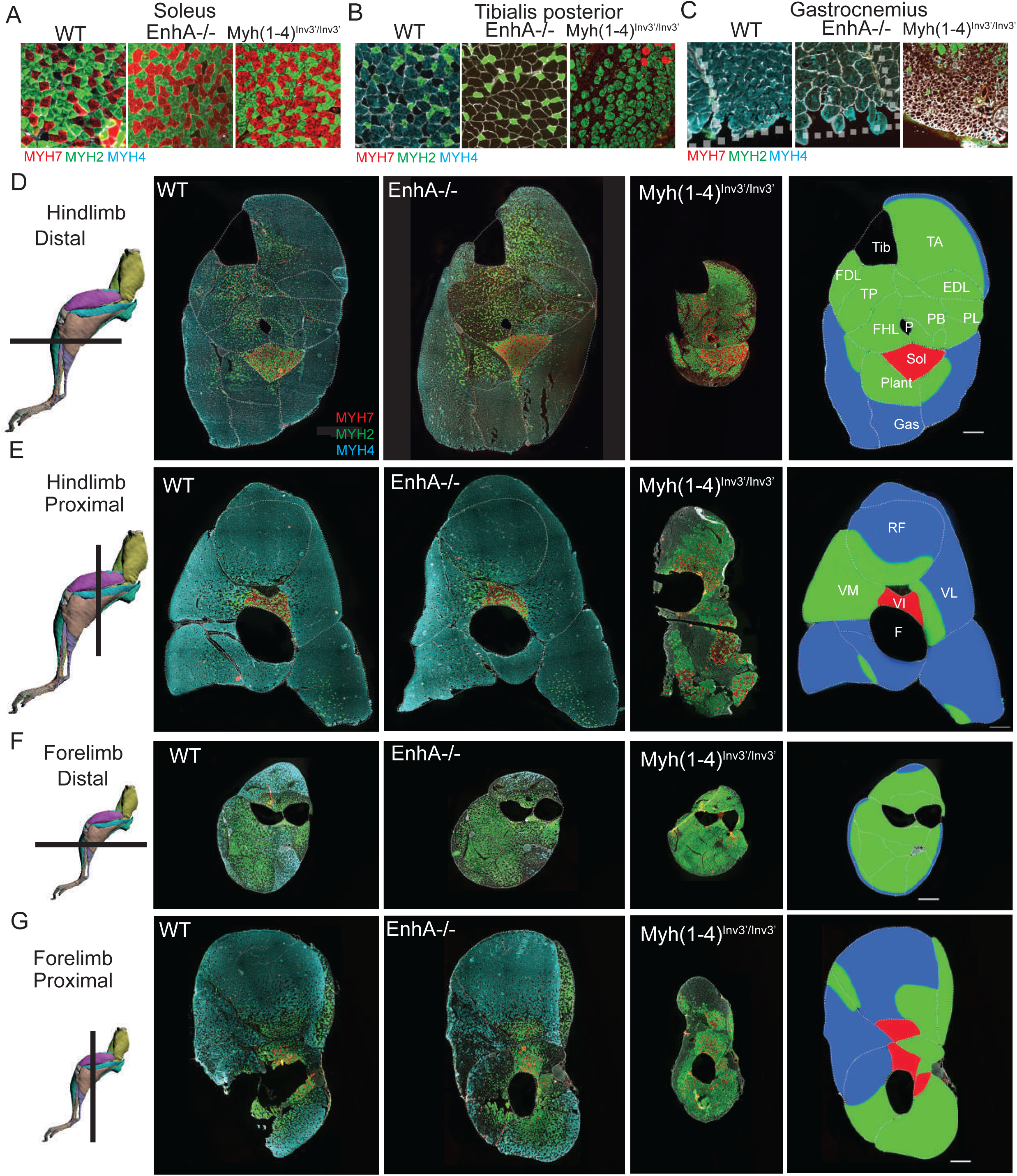
MYH expression established different groups of limb skeletal muscles in mutant animals. (A) Immunostaining against MYH7 (red), MYH2 (green) and MYH4 (blue) in soleus of WT, *EnhA^-/-^*, and *Myh(1-4)^Inv3’/Inv3’^*adult mice. The soleus is not affected in these mutant mice. (B) Immunostaining against MYH7 (red), MYH2 (green) and MYH4 (blue) in Tibialis posterior of WT, *EnhA^-/-^*, and *Myh(1-4)^inv3’/inv3’^*adult mice. In this muscle, the expression of MYH4 is lost in *EnhA^-/-^*, and in *Myh(1-4)^inv3’/inv3’^*. In this muscle an upregulation of MYH1 is observed in *EnhA^-/-^* mice and an upregulation of MYH2 in *Myh(1-4)^Inv3’/Inv3’^* mice. (C) Immunostaining against MYH7 (red), MYH2 (green) and MYH4 (blue) in peripheral Gastrocnemius of WT, *EnhA^-/-^*, and *Myh(1-4) ^Inv3’/Inv3’^*adult mice. The peripheral fibers of *Myh(1-4)^Inv3’/Inv3’^* Gas present a severe atrophy. In contrast these fibers are not affected in *EnhA^-/-^* mouse. (D) Comparison of the phenotypes of adult muscles in *EnhA^-/-^* and in *Myh(1-4) ^Inv3’/Inv3’^*allowed classification of distal hindlimb muscles in three major categories. The first group shown in red corresponded to the soleus that is not affected. The second group shown in green corresponded to muscles affected in *EnhA^-/-^* and *Myh(1-4)^Inv3’/Inv3’^* mutants. The third group shown in blue corresponded to muscles strongly affected in *Myh(1-4)^Inv3’/Inv3’^* but not in *EnhA^-/-^* mutants. (E) Same as (D) in the proximal part of the hindlimb. (F) Same as (D) in the distal part of the forelimb. (G) Same as (D) in the proximal part of the forelimb. For D, scale bar: 100μm. D-G: drawings of hindlimbs and forelimbs are from Charles et al ^41^.

The category of muscles similar to the Tibialis posterior (shown in green) was located between these two groups (Figure 6D-G). In the proximal part of the hindlimb, the group of muscles shown in blue was the most important and were severely affected in *Myh(1-4)^Inv3’/Inv3’^* mutants, whereas the same group of muscles was almost not affected in *EnhA^-/-^* mutants (Figure 6E). In the distal part of the forelimb, the group of muscles shown in green prevailed over the other (Figure 6F) whereas in the proximal part, the distribution of these muscles groups was more heterogeneous (Figure 6G). Altogether these results revealed that limb skeletal muscles could be classified in 3 major categories with distinct properties and possessing different codes of transcription factors controlling their plasticity.

## Discussion

In adult muscles the contraction and general metabolic properties of the specialized myofibers are dictated by the expression of specific slow MYH7 and fMYH subtypes (MYH2, MYH1, MYH4, MYH13) ^4, 5^. Transient transfection experiments of GFP reporters previously suggested that the proximal (800-1000bp) promoters of the *Myh2*, *Myh1* and *Myh4* genes were sufficient to drive their spatial expression in adult muscles ^42^. By combining single-nucleus ATAC-seq, ChIP-seq and 4C-seq data from adult fast and slow skeletal muscles, we show here that *fMyh* genes, with the exception of *Myh3*, are regulated by a shared super enhancer. In fast-type myonuclei this SE interacts dynamically with the activated promoters of the locus by 3D chromatin looping. By using rainbow transgenic mouse models of the locus and knock-out mouse models of the SE, we show that this SE controls the level and the spatio-temporal specificity of f*Myh* genes expression in myonuclei and myofibers through exclusive interactions with their promoters. By disrupting the organization of the f*Myh* locus, we uncover positional heterogeneity within limb skeletal muscles that may underlie selective muscle vulnerability observed in certain human neuromuscular diseases.

### Properties of the f*Myh*-SE: competition of f*Myh* promoters for the shared SE

SEs, composed of multiple enhancers, allow a more efficient recruitment of coactivators than conventional enhancers. During this process, multimolecular assemblies form by phase separation, allowing aggregation of the transcriptional machinery ^16, 23^. Known SEs have been described to achieve a relatively constant high transcriptional activity, contrasting with the transcriptional bursts provided by typical enhancers that lead to episodic gene expression ^15, 43^. We show here that the fine spatio-temporal expression of the f*Myh* genes is governed by a SE, which interacts with f*Myh* promoters by 3D chromatin looping, and is engaged in exclusive interactions with a single promoter at the locus. RNAscope experiments with f*Myh* premRNA probes demonstrated previously the coordinated firing of both alleles of specific f*Myh* genes in adult myonuclei ^10^, suggesting that specific SE-promoter loops may form simultaneously on both alleles of a given f*Myh* gene in the majority of body nuclei of each adult myofiber. Altogether, these results show that the f*Myh*-SE activates a single gene at the f*Myh* locus, suggesting the f*Myh*-SE cannot simultaneously activate two promoters and arguing against flip-flop mechanisms between the f*Myh*-SE and the different promoters of the locus ^17, 44, 45^. Whether this apparently non-stochastic gene expression in adult myofibers is true for all muscle genes governed by the SE remains to be established. At the f*Myh* locus, we suspect that these exclusive interactions between the SE and specific promoters are responsible for the high level of f*Myh* expression and to prevent the expression of two different MYH in adult myofibers. To test if these exclusive interactions result from a competition between the SE and the associated f*Myh* promoters, we analyzed the consequences of *Myh1* and *Myh4* deletion. Muscles of adult *Myh(1-4)^Del/Del^* mutant were composed of myofibers expressing MYH2, MYH8 or MYH13. Interestingly, *Myh2,* which is closest to the SE, was upregulated only in the deep regions of skeletal muscles, while in more peripheral myofibers where *Myh4* is normally predominantly expressed *Myh8* or *Myh13* were activated. These results suggested that *Myh2* cannot be activated in these peripheral myofibers, even in absence of *Myh1* and *Myh4,* and that competition between the promoters varies depending on the muscle position inside the limb, probably due to the differential enrichment of specific transcription factors in deep and peripheral muscles. These experiments demonstrated that some myofibers have the ability to switch from one specific promoter to another non-random promoter, suggesting that the transcription factors bound to *Myh8* and *Myh13* promoters in adult WT limb myofibers are able to interact with the SE, but compete less efficiently than those bound to *Myh4* probably due to a lower frequency of interactions. In adult wt limb muscles these preferential interactions concur to favor *Myh4* at the expense of *Myh8* and *Myh13* expression. Interestingly, even in *Myh(1-4)^Del/Del^* mutant, very few hybrid fibers were detected ^10, 34^, suggesting that most nuclei within each fiber activated a single gene, and that the SE was still contacting a single promoter in an exclusive manner. Such exclusive interactions were also detected at the *ß-Globin* locus where the LCR/SE interacts with a single promoter and where the order and the distance between the LCR and the *Globin* genes dictates their temporal expression ^19, 46^ but were not detected at *α-Globin* locus, where all promoters interact with the SE in a common nuclear compartment ^47^.

### The SE is composed of individual enhancer elements with redundant activities

To precise the role of the potential enhancer elements composing the SE, we focused on two elements, enhancer A and enhancer B. Deletion of enhancer A led to an upregulation of *Myh1* in myofibers of peroneal muscles, without up-regulation of the nearest *Myh2* promoter, again suggesting that the SE deleted of enhancer A still contacts a single promoter at a time with physically exclusive interactions. Down-regulation of *Myh4* in enhancer A mutant was observed only in certain muscles. This implies that the SE is composed of individual enhancer elements that may have redundant activities under the control of muscle identity genes, as in Drosophila ^48, 49^. This redundancy may contribute to the expression of a single gene at the locus. Altogether, we did not identify within the SE a specific enhancer responsible for driving the expression of either *Myh1*, *Myh2* or *Myh4* in all skeletal muscles. We cannot exclude that analysis of deletion of other snATAC-seq peaks may reveal a “master” enhancer element inside the SE. Alternatively the f*Myh*-SE may be composed of redundant elements, none of which absolutely required for driving efficient and specific gene expression, as fund at the *α-globin* locus and for enhancers controlling limb and digits morphogenesis where partially redundant enhancers are suspected to provide both flexibility and robustness of gene expression ^38, 50–53^.

### Importance of the f*Myh*-SE for human pathologies

Our experiments reveal the importance of the f*Myh*-SE for muscle integrity and function. Deletion of the SE induced impaired ability to breathe leading to death at birth. This deletion impaired *Myh1*, *Myh2*, *Myh4* and *Myh8* gene expression in skeletal muscles of E18.5 fetuses (one day before birth in C57BL/6N mouse strain), demonstrating the involvement of the SE to control their expression. Absence of *Myh8* expression may be involved in the death of the mutant animals. The requirement of *Myh8* expression during fetal development for efficient muscle contraction at birth is supported by the phenotype of *Myod^-/-^;Nfatc2^-/-^* mice, where *Myh8* is no more expressed in intercostal muscles. These mutant mice do not survive after birth due to their inability to breathe ^54^. In agreement we identified strong sarcomerisation defects associated with Actin aggregates in f*Myh*-SE^-/-^ E18.5 mutant myofibers at the limb and diaphragm level, suggesting a complete absence of MYH and their inability to contract. The SE is present as well in the human f*MYH* locus, and could be involved in the control of the f*MYH* genes as in mice. *MYH8* and *MYH2* are expressed during human fetal development, *MYH1* is detected after birth ^8, 55^, while *MYH4* is only expressed in extra ocular myofibers due to mutations in its promoter region ^36, 56^. Absence of *MYH2* is associated with early onset myopathy characterized by mild generalized muscle weakness with predominant involvement of muscles of the lower limbs, and by ophtalmoplegia ^57^. In contrast, *MYH1* mutations have not yet been reported and *MYH8* mutations do not seem to be associated with trismus- pseudocamptodactyly ^58^, contrarily to what was previously suspected. Mutations or deletions of the f*MYH*-SE have not yet been identified in human pathologies. Congenital myopathies can be associated with Actin aggregates, fiber type disproportion or arthrogryposis ^59–61^, but not all of these myopathies have been characterized at the genetic level.

The human body is composed of more than 600 different skeletal muscles, each with specific functions and properties. This heterogeneity is reflected in the spectrum of clinical manifestations of neuromuscular diseases where some specific muscles are affected while other are spared, depending on the pathology ^1^. Whole-body magnetic resonance imaging and muscle ultrasound in patients affected by Collagen VI deficiency, Dystrophin deficiency or in ALS showed that specific muscles or specific group of myofibers inside a muscle mass can be specifically affected, while others are spared ^62–64^. Little is known about the mechanisms driving this variability in susceptibility and understanding the underpinning mechanisms is a major challenge to develop adapted targeted therapies. By disrupting the organization of f*Myh* at the locus, we uncovered positional heterogeneity within limb skeletal muscles and defined three major categories of limb muscles. These three categories of stereotyped muscles are differentially positioned in distal and proximal forelimbs and hindlimbs. We suspect that this heterogeneity may be the consequence of different genetic programs that lead to the activation of groups of genes associated with *Myh4*, or *Myh1* or *Myh2* expression. We and others ^10, 65, 66^ revealed recently an unsuspected genetic diversity of *Myh4*+ and other myofiber types with, in mouse hindlimbs, at least three subclasses of *Myh4* + myonuclei and several subclasses of *Myh1+* and *Myh2*+ myonuclei. Whether this diversity is at the origin of the deep/superficial gradient of muscle susceptibility observed in the present study and in certain neuromuscular diseases remains to be tested.

## Methods

### Animals

Animal experimentations were carried out in strict accordance with the European STE 123 and the French national charter on the Ethics of Animal Experimentation. Protocols were approved by the Ethical Committee of Animal Experiments of the Institut Cochin, CNRS UMR 8104, INSERM U1016. We used 6-8 weeks old C57BL/6N mouse female for most of our experiments. Mice were anesthetized with intraperitoneal injections of ketamine and xylazine and with subcutaneous buprecare injections before denervation was performed by sectioning of the sciatic nerve in one leg. All efforts were made to minimize animal suffering, and to reduce the number of animals required for the experiments.

### BAC targeting constructs and *Myh* locus modifications

For the construction of the targeting vector pGEM-T-EasyMyh2YFP, C57BL/6N mouse DNA was first used as a template to clone 5’ arm and 3’arm of *Myh2* with forward 5’- GAA TGA TTT CAT TGC TAC TTC -3’ and reverse HindIII 5’- GCT CAT GAC TGC TGA ACT CAC -3’, and forward HindIII 5’- AGT CCG AAA AGG AGC GAA TC -3’ and reverse 5’- GGT GAC TTC TAG TGA CTG AG -3’, respectively. The 5’ arm and 3’arm fragments were cloned into a pGEM-T-Easy vector with HindIII in-between to make pGEM-T-Easy*Myh2*. The Yellow Fluorescent Protein (YFP) coding sequence was PCR amplified (PHUSION, Thermofisher) and cloned in pBluescriptSK+ using EagI-XbaI sites provided by the primers. Fragments containing three polyA sequences (rabbit β-globin, HSV-TK, and BGH) and LoxP-kanamycin-LoxP were then extracted from preexisting constructs and introduced downstream of YFP. The whole YFP- 3pA-LoxP-kana-LoxP fragment was amplified (PHUSION, Thermofisher) with forward 5’- CAG CAG TCA TGA GCA TGG TGA GCA AGG GCG AGG AG-3’ and reverse 5’- CTC CTT TTC GGA CTA CGA CTC ACT ATA GGG CGA ATT G-3’ primers. The resulting amplicon features 15bp homology in 5’ and 3’ extremities with the targeting arms allowing Sequence and Ligation Independant Cloning (SLIC) into the HindIII digested pGEM-T- Easy*Myh2* plasmid (GeneArt Seamless Cloning and Assembling kit, Thermofisher).

Similarly, for the construction of the targeting vector pGEM-T-Easy*Myh1*Tomato, targeting arms were PCR generated from C57BL/6N mouse DNA and assembled together with HindIII in-between (pGEM-T-Easy-*Myh1* : 5’ arm forward 5’- CAT CCA GCA TGT GTT CTC AGA GGT -3’, reverse HindIII 5’- ACT CAT GGC TGC GGG CTA TT -3’ ; 3’arm forward HindIII 5’- GTC TGA AAA GGA GCG AAT CGA G -3’, reverse 5’- AGT AGG TCT GCA TCA AGA GAG GG -3’). The PCR amplified tandem-dimer-Tomato (TdTomato) coding sequence was cloned in Bsp120I-XbaI of pBluescriptSK+. The three polyA signals and Lox2272-kanamycin- Lox2272 cassettes were subsequently added downstream of TdTomato. For SLIC, 5’- CCG CAG CCA TGA GTA TGG TGA GCA AGG GCG AGG AG -3’ and 5’- GCT CCT TTT CAG ACA CGA CTC ACT ATA GGG CGA ATT G -3’ primers were used and pGEM-T-Easy- Myh1 linearized with HindIII. The targeting vector pGEM-T-EasyMyh4CFP was generated by SLIC of a CFP-3pA-LoxN-KanamycinLoxN PCR fragment into HindIII linearised pGEM-T- EasyMyh4 (C57BL/6N mouse DNA targeting arms : 5’arm forward 5’- CCC AAG TGC TGG AAT TGA AAG TGT -3’, reverse HindIII 5’- ACT CAT GGC TGC GGG CTA TT -3’ ; 3’arm forward HindIII 5’- GTC TGA AAA GGA GCG AAT CG -3’, reverse 5’- GCT AAC TAT CAG CAC GTG CA -3’) using forward 5’- CCG CAG CCA TGA GTA TGG TGA GCA AGG GCG AGG AG -3’ and reverse 5’- GCT CCT TTT CAG ACA CGA CTC ACT ATA GGG CGA ATT G -3’ primers.

A 222kb Bacterial Artificial Chromosome (BAC) from a C57BL/6J mice genomic library ^67^ containing the whole *Mhy2* to *Mhy4* locus surrounded by 80kb of genomic DNA upstream and 46kb downstream is chosen (RP23-61C14 ; CHORI BACPAC resources) to carry out genetic alterations using λ-red recombination ^68^. To remove Lox motifs preexisting on the pBACe3.6 backbone which will later interfere with our strategy of recombination, BAC DNA amplified in DH10b is extracted (Nucleobond MIDI XTRA, Macherey-Nagel), checked by Acc65I-NotI complex restriction profile, and transformed by electroporation into SW105 competent cells. BAC DNA from several transformants is extracted and checked using the same complex restriction profile against the parental one. Removal of LoxP is carried out on one bacterial clone made competent then induced for recombinase expression by 15 minutes incubation at 42°C by electroporation of a 1.85kb BamHI-NotI DNA fragment purified from pTamp- BACe3.6 (gift of Dr V. Besson) conferring ampicillin resistance. BAC DNA from recombinant ampicillin resistant clones is extracted and checked against parental DNA using Acc65-NotI or MfeI-NotI complex restriction profiling. Similarly, removal of Lox511 is performed on one ampicillin resistant clone using a 2.2kb KpnI-BamHI fragment purified from pSKTHygroBACe3.6Lox511 (gift of Dr J. Hadchouel) which confers hygromycin resistance to recombinant clones. DNA from one clone is then transformed into SW106 cells harboring Cre-inducible expression under arabinose treatment ^69^ for further targeting step.

Sequential *Myh2*, *Myh4* and *Myh1* locus modifications are performed by 3 rounds of competent bacterial clone electroporation using a 3.75kb NotI transgene purified from each respective pGEMTe-based targeting vector described above followed by kanamycin selection of recombinant clones, BAC DNA extraction, complex restriction profiling against parental DNA, then from a proper recombinant clone floxing-out kanamycin resistance by 0.1% arabinose treatment, BAC DNA extraction and again complex restriction profiling against parental DNA. Enzymes combinations are as follows: KpnI+NotI and MfeI+NotI for *Myh2*-YFP, *Myh4*-CFP- kana and *Myh1*-TdT; MfeI+NotI and BamHI+NotI for *Myh4*-CFP. The final transgenic BAC DNA is then transferred back to DH10b cells for better extraction yield (Nucleobond BAC100, Macherey-Nagel). DNA is resuspended in 10mM Tris-HCL pH 7.0, 1mM EDTA, 100mM NaCl. The final transgenic BAC DNA is then transferred back to DH10b cells for better extraction yield (Nucleobond BAC100, Macherey-Nagel). DNA is resuspended in injection buffer (10mM Tris-HCL pH 7.0, 1mM EDTA, 100mM NaCl), and 200ng filtrated through drop dialysis against the filtration buffer for 1h using Millipore cellulose ester disc membranes VMWP 0.05µm (Ref# VMWP02500).

### Immunohistochemistry

Immunostaining against YFP and MYH2 were performed on soleus and immunostaining against Tomato and Myh1 were performed on quadriceps. Muscles were fixed 30 minutes in PFA 2% with 0,2% Triton at 4°C. After overnight 10% sucrose treatment, muscles were embedded with TissuTEK OCT (Sakura) and frozen in cold isopentane cooled in liquid nitrogen. For immunostaining against MYH4, MYH2, MYH7, and Laminin, adult legs without fixation and without skin were embedded with TissuTEK OCT and directly frozen in cold isopentane cooled in liquid nitrogen Muscles were conserved at -80°C and cut with Leica cryostat 3050s with a thickness of 10µm. Cryostat sections were washed 3 times 5 minutes with PBS and then incubated with blocking solution (PBS and 10% goat serum) 30 minutes at room temperature. Sections were incubated overnight with primary antibody solution at +4°C, then washed 3 times for 5 minutes with PBS and incubated with secondary antibody solution 1 hour at room temperature. Sections were further washed 3 times 5 minutes and mounted with mowiol solution and a glass coverslip. Images were collected with an Olympus BX63F microscope and a Hamamatsu ORCA-Flash 4.0 camera. Images were analyzed with ImageJ program. The references of the antibodies used are listed in Table S1.

### RNA extraction and quantification

RNA extractions from adult skeletal muscles were performed using TRIzol reagent (ThermoFischer) following the manufacturer’s protocol. Muscles were lysed with Tissue lyser (Quiagen) in TRIzol solution. RNA was precipitated with isopropanol. cDNA synthesis was performed with Superscript III kit (Invitrogen) using 1µg of RNA. RT-qPCR were performed using Light Cycler 480 (Roche) with the Light Cycler 480 SYBR Green I Master Kit (Roche) following the manufacturer’s protocol with 40 cycles at 95°C for 15 seconds, 60°C for 15 seconds, and 72°C for 15 seconds. We used 36B4 housekeeping gene to normalize the expression level between different samples. The sequences of the oligonucleotides used are listed in Table S2.

### Single nucleus ATAC-seq from skeletal muscle

We use the 10X genomic nuclei Isolation for Single Cell ATAC Sequencing protocol (CG000169 | Rev B) with some changes. 12 quadriceps and 12 soleus were dissected and pulled in cold PBS. PBS was removed and muscles were minced 2 minutes in 1 ml of cold ATAC-lysis buffer (10mM Tris-HCl pH7.4, 10mM NaCl, 3mM MgCl2, 1% BSA and 0.1% Tween-20 in Nuclease-Free Water). 6ml of cold ATAC-lysis buffer were added and muscles were lysed on ice. After 3 minutes the lysate was dounced with 10 strokes of loose pestle avoiding too much pressure and air bubbles. After douncing, 8 ml of wash buffer were added and the homogenate was filtered with 70μm, 40μm and 20μm cell strainers. Nuclei were pelleted by centrifugation for 5min at 500g at +4°C. Next, we used the Chromium Single Cell ATAC kit according to the manufacturer’s protocol. Nuclei were resuspended in nuclei buffer from the kit, transposed 1 hour at 37°C. We loaded around 6000 nuclei into the 10X Chromium Chip. GEM incubation and amplification were performed in a thermal cycler: 72°C for 5 min, 98°C for 30 sec and 12 repeated cycles of 98°C for 10 sec, 59°C for 30 sec and 72°C for 1 min. Post GEM Cleanup using DynaBeads MyOne Silane Beads was followed by library construction (98°C for 45 sec, cycled 12 x 98°C for 20 sec, 67°C for 30 sec, 72°C for1 min). The libraries were constructed by adding sample index PCR, and SPRIselect size selection. The fragment size estimation of the resulting libraries was assessed with High SensitivityTM HS DNA kit runed on 2100 Bioanalyzer (Agilent) and quantified using the QubitTM dsDNA High Sensitivity HS assay (ThermoFisher Scientific). Libraries were then sequenced by pair with a HighOutput flowcel using an Illumina Nextseq 500.

### Single-nucleus ATAC-seq analysis

A minimum of 10 000 reads per nucleus were sequenced and analyzed with Cell Ranger Single Cell Software Suite 3.0.2 by 10X Genomics. Raw base call files from the Nextseq 500 were demultiplexed with the cellranger-atac mkfastq pipeline into library specific FASTQ files. The FASTQ files for each library were then processed independently with the cellranger count pipeline. This pipeline used STAR21 to align reads to the Mus musculus genome. Once aligned, barcodes associated with these reads –cell identifiers and Unique Molecular Identifiers (UMIs), underwent filtering and correction. The subsequent visualizations, clustering and differential expression tests were performed in R (v 3.4.3) using Seurat36 (v3.0.2) ^70^, Signac (v0.2.4) (https://github.com/timoast/signac) and Chromvar (v1.1.1) ^71^. Quality control on aligned and counted reads was performed keeping cells with peak_region_fragments > 3000 reads and < 100000, pct reads in peaks > 15, blacklist ratio < 0.025, nucleosome_signal < 10 and TSS.enrichment > 2. We get 6037 nuclei in total and we detected 132 966 peaks (Figure S1). The motif activity score was analyzed by running chromVAR (Figure 3C-E).

### ChIP-seq analysis

Fastq files of quadriceps femoris and soleus H3K27ac ChIP-seq ^24^ were download from the GEO database (accession number GSE123879). The reads were aligned to the mouse mm10 genome using bowtie2 ^72^ and peaks were called by MACS2 ^73^ using q value cutoff = 0.05. ROSE algorithm ^14^ was applied to identify and rank the enhancers based on H3K27ac ChIP-seq signal, with a stitching distance of 12.5 kb.

### Nuclei purification from adult skeletal muscle for 4C-seq

Nuclei purification from adult skeletal muscle has been performed as previously described ^74^ with some modifications. After dissection, 16 soleus or 8 quadriceps were resuspended in 1mL of hypotonic buffer (25mM Hepes-KOH pH 7.8, 10mM KCL, 1.5mM MgCl2, 0.1% NP40, PIC 1X (complete protease inhibitor Roche), PMSF 1mM) in a 2ml tube for 5 min at +4°C. Muscles were sliced with a scissor for 30 sec. The small pieces of muscles were transferred into a round tube of 15mL at +4°C and 4ml of cold hypotonic buffer was added. After 5 min the solution was homogenized for 15 seconds with an Ultra-Turrax (IKA) at a speed of 17,500 rpm. The solution was transferred in a 15 ml Falcon tube and crosslinked with 2% formaldehyde (in a volume of 10ml of hypotonic buffer) at room temperature during 10 min. 1,43 ml of cold glycine (1M) was added to quench the formaldehyde for 5 min at +4°C while shaking. The crosslinked nuclei were dounced 10 times with a loose pestle and then centrifuged at 1000 Rcf for 10 min at +4°C. The nuclear pellet was resuspended in 5ml of hypotonic buffer and filtered with 70µm and 40µm cells strainers. The nuclei were pelleted with centrifugation at 1000 rcf for 10 min, snap frozen into liquid nitrogen and stored at -80°C.

### 4C-seq

4C-seq experiments have been performed as previously described ^75^ with some modifications. Purified crosslinked nuclei from 160 soleus and 80 quadriceps were pooled together to have 10^7^ nuclei per conditions. PCR primers were designed for each viewpoint according to the protocol. The first digestion was done with DpnII (New England Biolabs) and the second with NlaIII (New England Biolabs). For each viewpoint 800ng of 4C template was amplified by PCR. The samples were sequenced on the Illumina NextSeq 500 platform, using 75 bp single end reafs. The analysis of the data has been done using the HTSstation 4C-seq pipeline ^76^. Briefly, the sequences were demultiplexed, then aligned to the reference genome (mm10) and normalized. Hi-C data in mouse ES cells were obtained from the 3D Genome Browser website (http://promoter.bx.psu.edu/hi-c/view.php). ChIP-seq data against CTCF in mouse ES cells and DNase I hypersensitive site in adult fast skeletal muscle were obtained from the ENCODE database. The sequences of the oligonucleotides used for 4C-seq are listed in Table S3.

### Mouse generation by CRISPR/Cas9

SgRNA and Cas9 purified protein were produced by the TACGENE platform. The SgRNA were designed with the Crispor program (http://crispor.tefor.net/) ^77^. SgRNA are produced by T7 Hiscribe transcription kit (New England Biolabs) and purified by EZNA microelute RNA clean up kit (Omega biotek). The DNA used for transcription was produced by overlapping PCR. For each cut sites, 3 different sgRNA were designed and tested *in vitro* by transfection in MEF cells. The deletions were performed by injecting into oocytes between 1 and 5 pg of sgRNA (60ng/μl) cutting at both sides of the deletion and of the Cas9 protein (30 μM). Oocytes where reimplanted into a pseudopregnant females. Mutant mice were screened by PCR and confirmed by sequencing. The list of the sgRNA and PCR primers used for screening are listed in Table S4.

### FISH with amplification (RNAscope) on isolated fibers

RNAscope® Multiplex Fluorescent Assay V2 was used to visualize fast *Myh* pre-mRNAs and mRNAs. Twenty different pairs of probes against the first intron of each fast *Myh* transcript were designed by ACDbio. Muscles were dissected and immediately fixed in 4% PFA at +4°C during 30 minutes. After fixation muscles were washed 3 times in PBS for 5 min. Myofibers were dissociated mechanically with small tweezers and fixed onto Superfrost plus slides (Thermo Fischer) coated with Cell-Tak (Corning) by dehydration at +55°C during 5 min. Slides were then proceeded according to the manufacturer’s protocol: ethanol dehydration, 10 min of H2O2 treatment and 30 min of protease IV treatment. After hybridization and revelation, the fibers were mounted under a glass coverslip with Prolong Gold Antifade Mountant (Thermofischer). Myofibers were imaged with a Leica DMI6000 confocal microscope composed by an Okogawa CSU-X1M1 spinning disk and a CoolSnap HQ2 Photometrics camera. Images were analyzed with Fiji Cell counter program.

### Statistical analysis

The graphs represent mean values ± SEM. Significant differences between mean values were evaluated using two-way ANOVA for Fig 2H, one way ANOVA with multiple comparisons for Fig 3D, Fig 4E, Fig 5C, Fig 5D, Fig 5E, Sup 8E, Sup 9F, Sup 9G, Sup 9H and student t test for Fig. 1B, Fig 2K with Graphpad 8.4.3 software.

### GEO data accession number

4C-seq data have been deposited in the NCBI Gene Expression Omnibus database (https://www.ncbi.nlm.nih.gov/geo/): accession number GSE168074

The reviewers can have (anonymous) access to the data using this link https://www.ncbi.nlm.nih.gov/geo/query/acc.cgi?acc=GSE168074 And using the password odklgemslhuvjgx

## Acknowledgments

We thank D.Duboule, S.Schiaffino, H.Amthor, M.Schuelke and F.Spitz for helpful discussions. We thank U.Schibler, S.Gautron and F.Britto for helpful critical reading of the manuscript. We thank T. Guilbert and F. Letourneur at the Cochin IMAG’IC and GENOM’IC platform for helpful advice. We acknowledge the High-throughput sequencing facility of the I2BC (Gif-sur-Yvette, France) for its sequencing and bioinformatics expertise. M.D.S was supported successively by a PhD fellowship from the Association Française contre les Myopathies (AFM), by a post doc fellowship from EUR-Gene and from the Agence Nationale pour la Recherche (ANR Myolinc R17062KK). Financial support to this work was provided by the AFM (n°16427, 21711 and 23012), the ANR (Myocodes R09108KK and Myolinc R17062KK), the Institut National de la Santé et de la Recherche Médicale (INSERM) and the Centre National de la Recherche Scientifique (CNRS) to P.M, by AFM (n° 21711) to J.D, and by AFM (n°19507 and 22946 Translamuscle) to F.R. Designed experiments, M.D.S, D.N, I.S, F.A, P.M. Performed experiments, M.D.S, I.S, S.B, M.Wu, F.A, R.P, F.L, M.D.C, A.S, J-P.C. Interpreted data, M.D.S, I.S, D.N, A.S, P.M. Bioinformatic analysis, M.Wo, M.D.S. Wrote the manuscript, M.D.S, P.M. with input from D.N, A.S, and J.D. Funding Acquisition, P.M, J.D, F.R.

---

**Figure S1.**
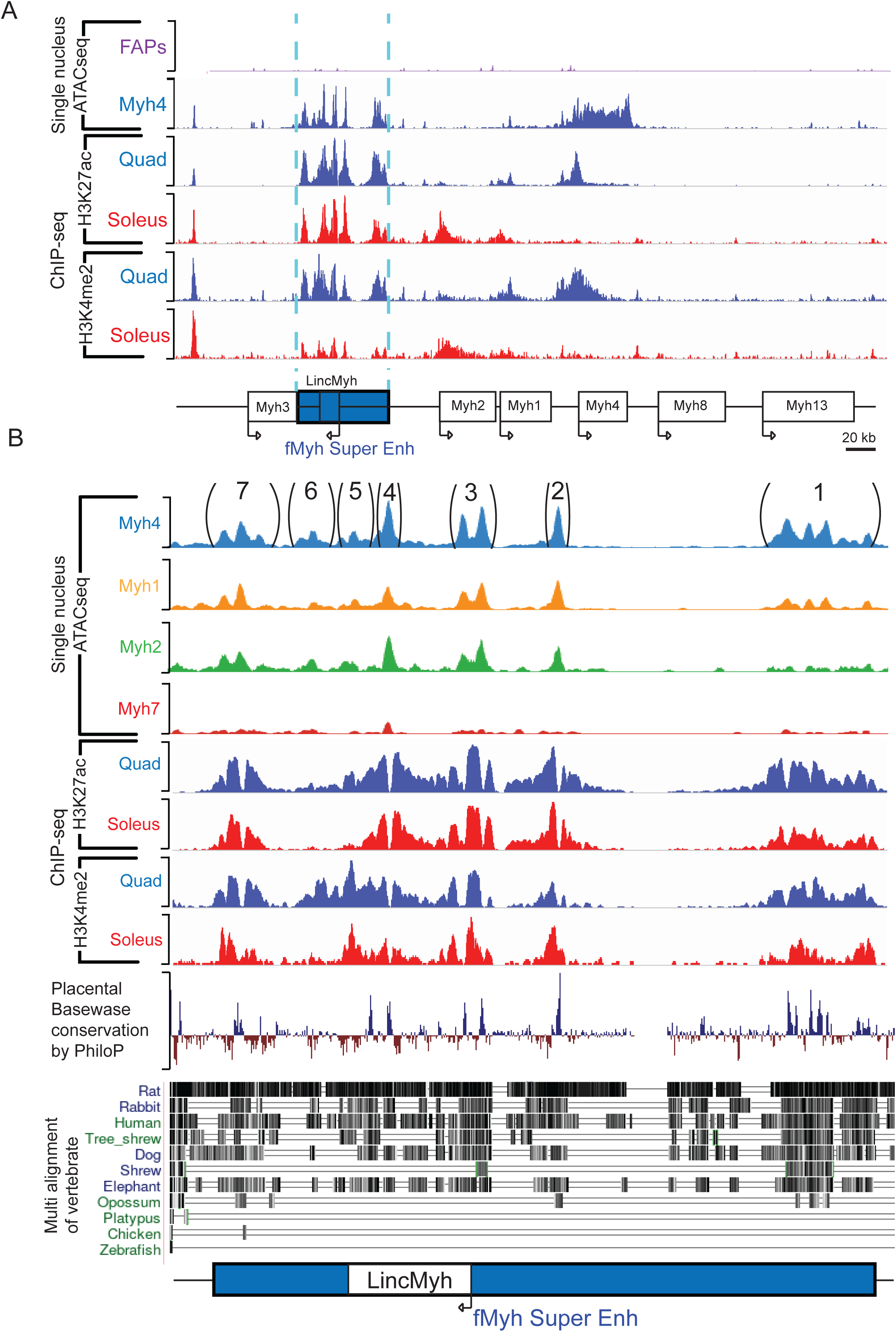
Active histone marks in slow and fast muscles. (A) Chromatin accessibility as identified in snATAC-seq experiments in FAPs and *Myh4* myonuclei, in ChIP-seq experiments ^24^ for H3K27Ac and H3K4me2 in quadriceps and soleus in the f*Myh* locus. (B) Same as (A) zoom in the f*Myh* SE, and its seven snATAC-seq peaks, placental base wase conservation by PhiloP and multi alignment of vertebrate DNA sequences of the 42kb SE.

**Figure S2.**
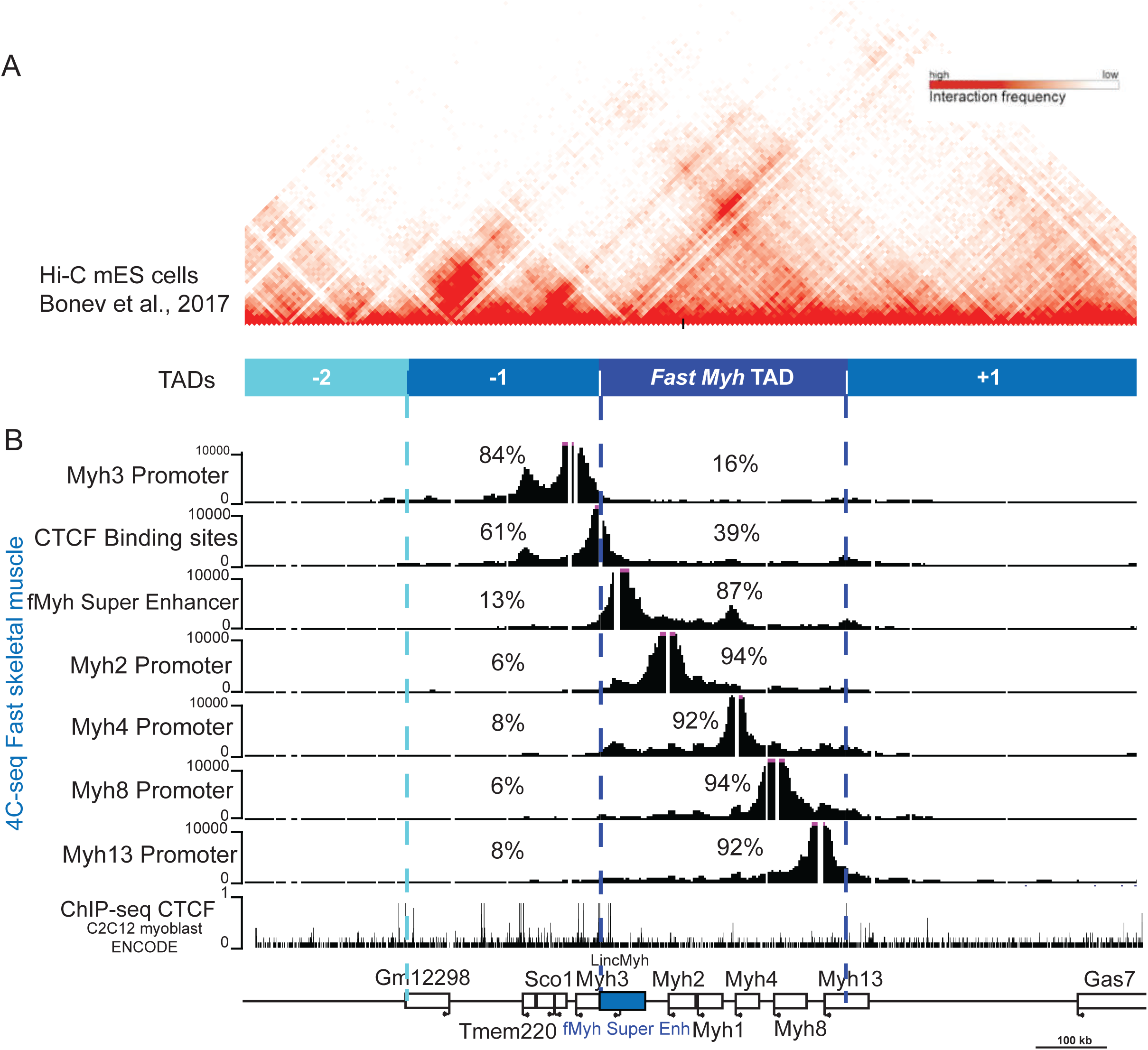
Distinct TADs partitioned the f*Myh* locus as determined by HiC-seq experiments. (A) Alignment of Hi-C data from mES cells ^32^. (B) 4C-seq data from adult fast skeletal muscles and ChIP-seq data against CTCF in C2C12 myoblast ^33^ in the f*Myh* locus and adjacent TADs. HiC-seq data revealed that the f*Myh* locus is organized in one TAD delimited by CTCF borders at the 3’ end of *Myh3* gene and at the 3’ end of *Myh13*, showing that *Myh3* does not belong to the same TAD that the other f*Myh* genes. The size of the TAD including *Myh1*, *Myh2* and *Myh4* is estimated at 350kb. For 4C-seq experiments, we used 7 distinct viewpoints in the f*Myh* locus to determine its organization in adult leg muscles.

**Figure S3.**
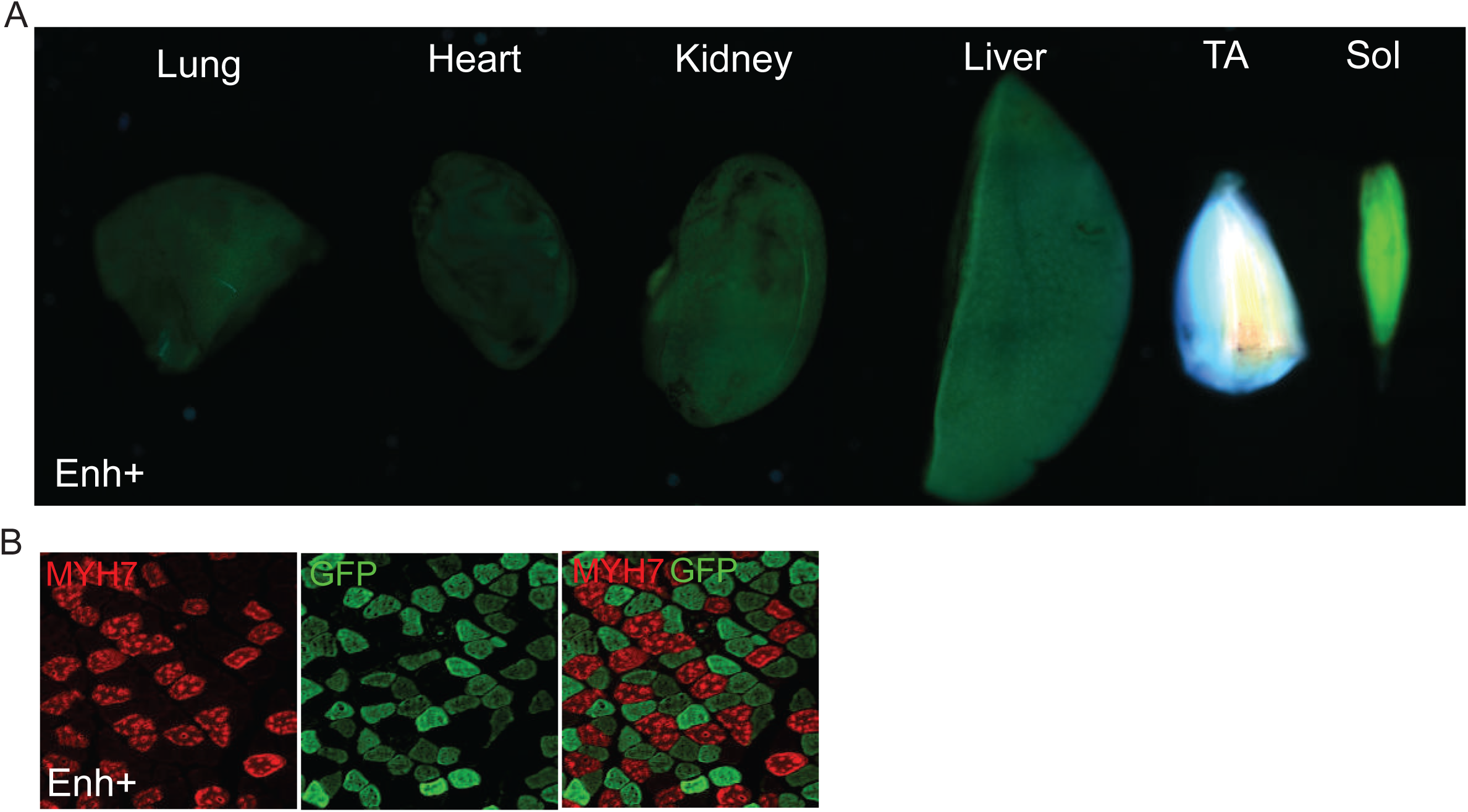
f*Myh* regulation in Enh+ transgenic animals. (A) Transgenes expression in several tissues in Enh+ line. The expression of YFP, Tomato and CFP is only detected in skeletal muscle (TA and soleus in the picture) and not in other organs like lung, heart, kidney, or liver. (B) Immunostaining against endogenous MYH7 (red) and YFP (with an anti GFP antibody in green) on adult soleus of Enh + mice showing that MYH7 fibers does not express YFP.

**Figure S4.**
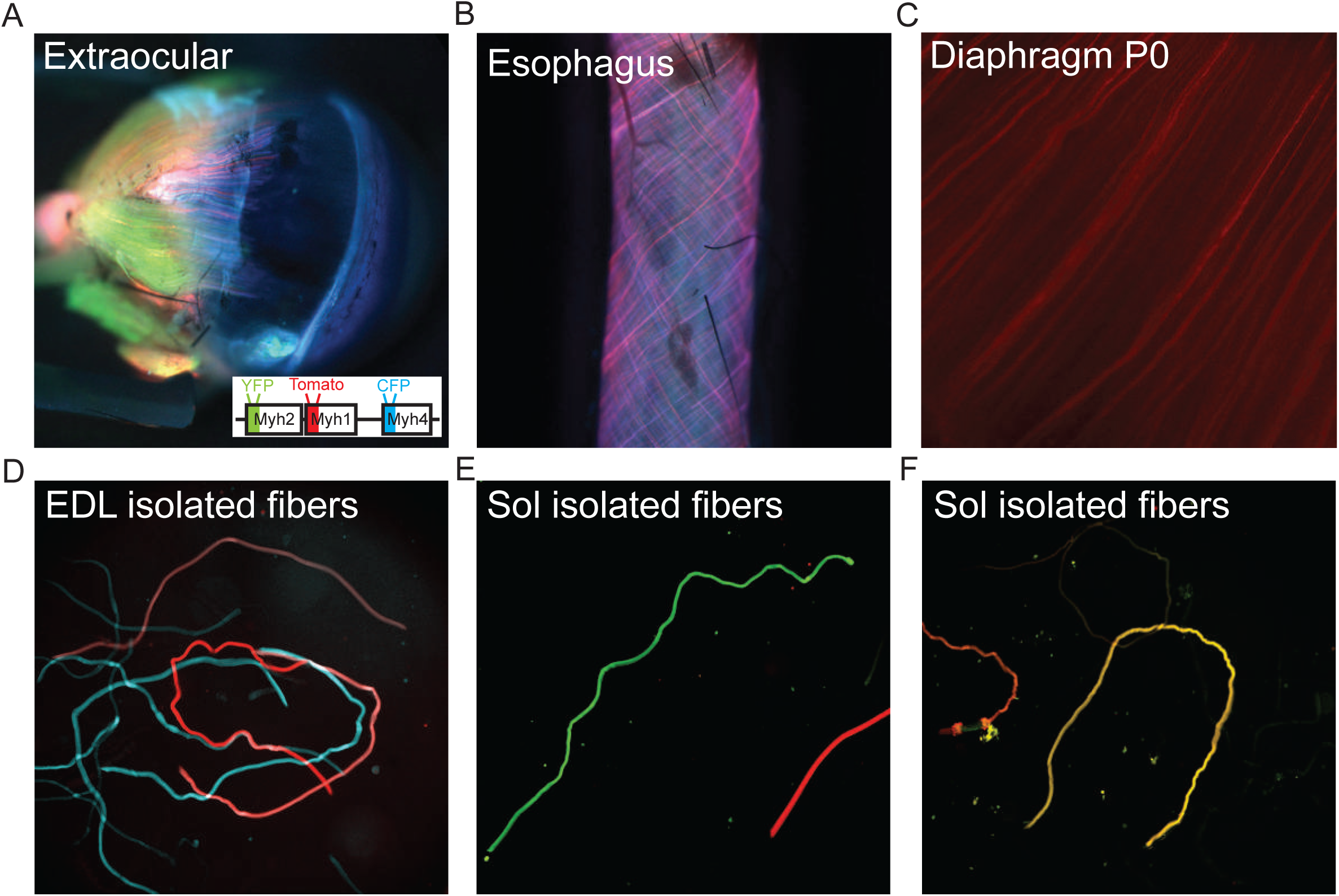
Expression of the transgenes in skeletal muscles of Enh+ mice. (A) Schema of the three inserted transgenes in the f*Myh* BAC, and expression of the transgenes in extraocular muscles. Red; Tomato, green; YFP and blue; CFP. (B) same as in (A) showing the esophagus muscle of an adult Enh+ mouse. (C) Same as (A) showing the diaphragm in P0 Enh+ mouse. (D) Transgenes expression in isolated fibers from the EDL. We could detect pure YFP, Tomato and CFP fibers and the expression of the transgene is similar all along the fibers. At the top, a pink Tomato-CFP hybrid fiber. (E) The majority of soleus fibers express only one transgene; Tomato or YFP. (F) A minority of fibers co-expressed Tomato and YFP in adult soleus muscle.

**Figure S5.**
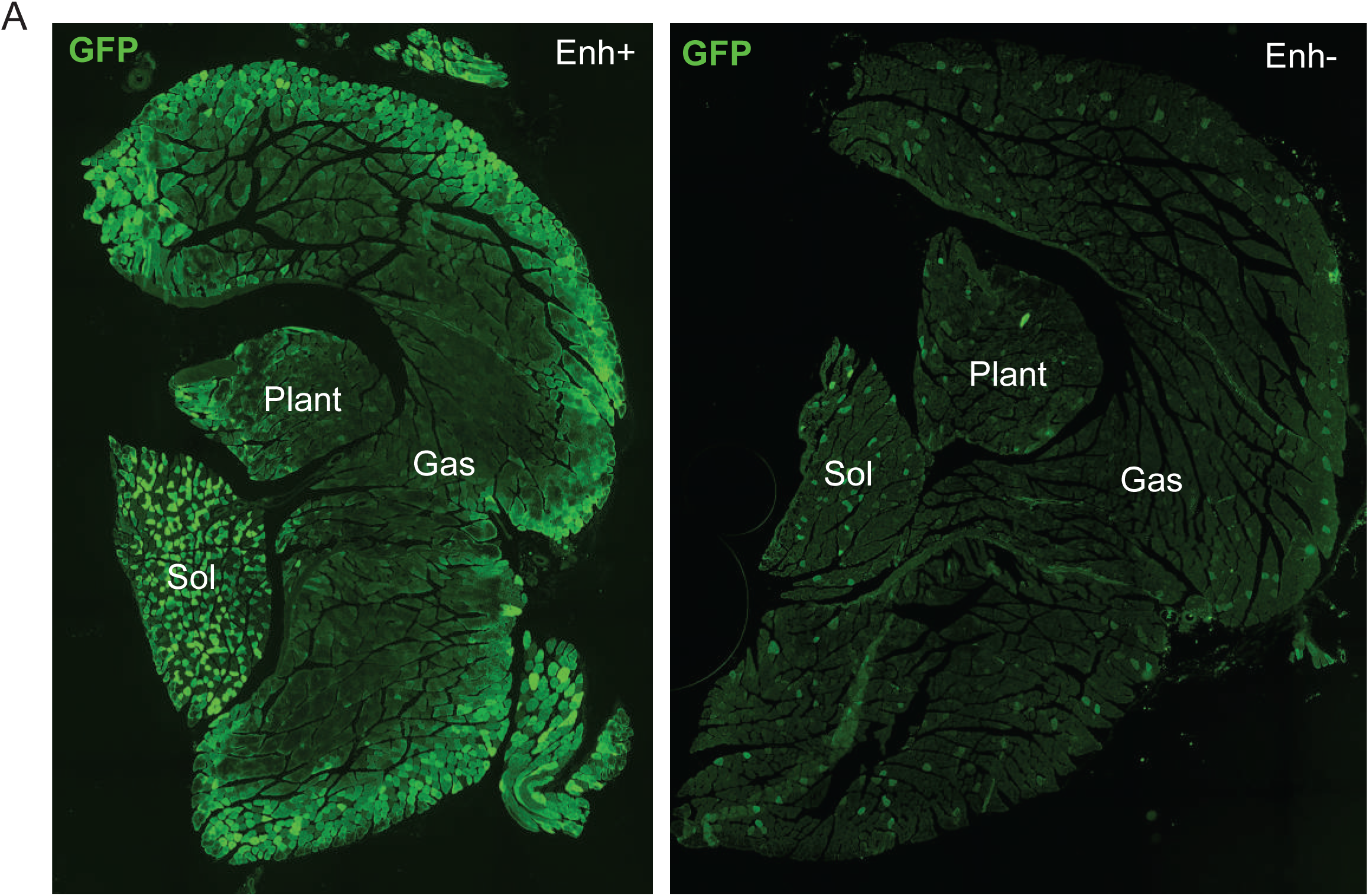
Transgene expression in Enh- mouse. (A) Immunostaining with GFP antibodies revealing YFP and CFP proteins on sections of adult soleus, plantaris and gastrocnemius in Enh+ (left) and Enh- (right) mice.

**Figure S6.**
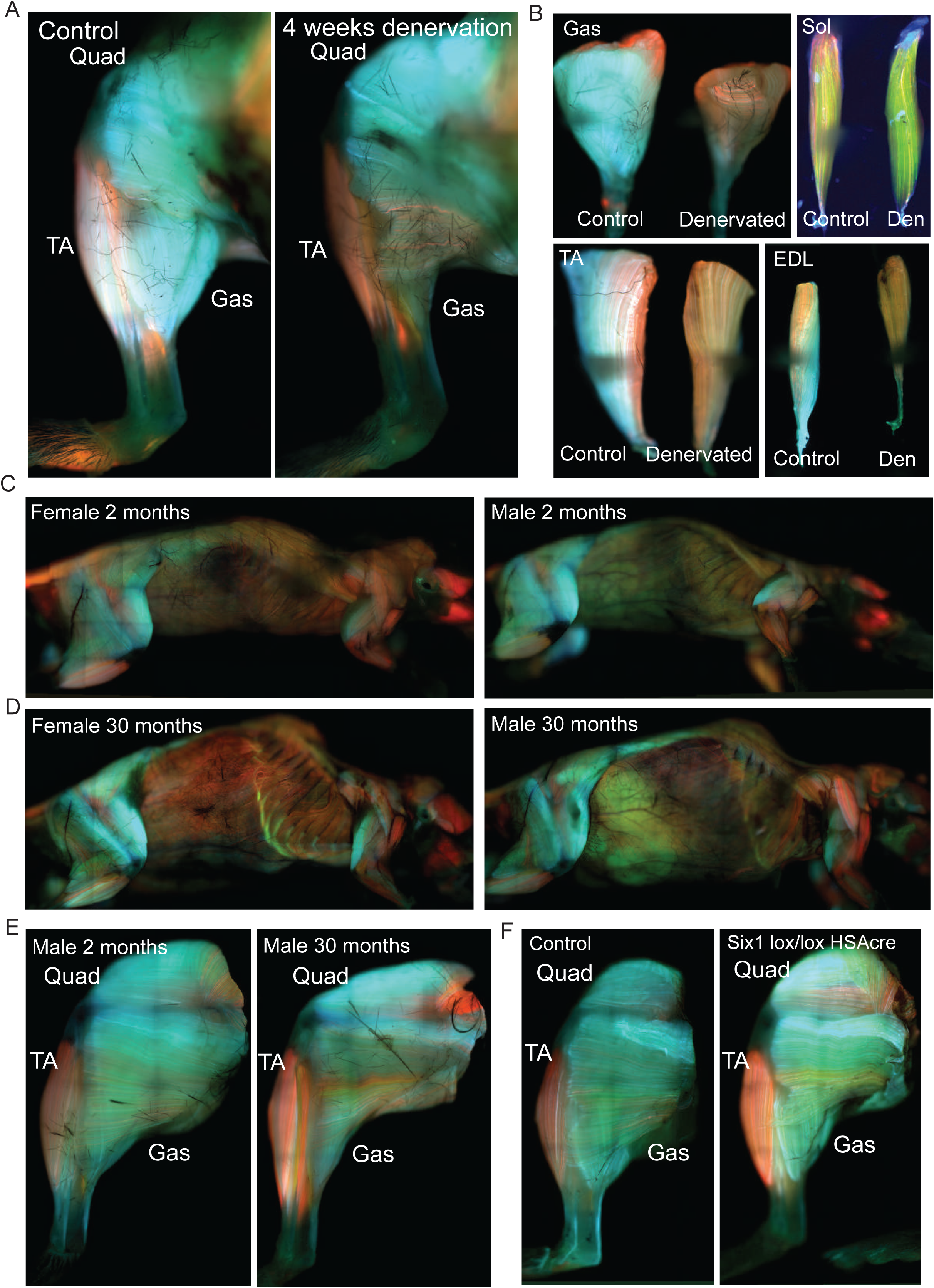
Transgenic Enh+ mice allowed identification of fiber-type modifications under various pathophysiological conditions. (A) Control and 4 weeks denervated hindlimb legs of Enh+ mice. Distal muscles are more affected by the absence of innervation than proximal muscles. A strong atrophy of the gastrocnemius is observed. (B) Same as (A), zoom in Gastrocnemius, Tibialis anterior, EDL and soleus. Denervation induced a fast to slow transition in these 3 muscles visible by the expression of the transgenes. (C) 2-month-old Enh+ female and male mice. Skeletal muscles of females have more Tomato/MYH1 fibers and less CFP/MYH4 fibers. (D) Same as (C) in 30- month-old Enh+ female and male. (E) hindlimb of 2- and 30-month-old Enh+ mice. (F) Control and *Six1^flox/flox^;HSA-CRE* hindlimb muscles of Enh+ mice. The absence of *Six1* in skeletal muscles induced a fast to slow fiber type switch. Gas: Gastrocnemius, TA: tibialis anterior, Sol: soleus.

**Figure S7.**
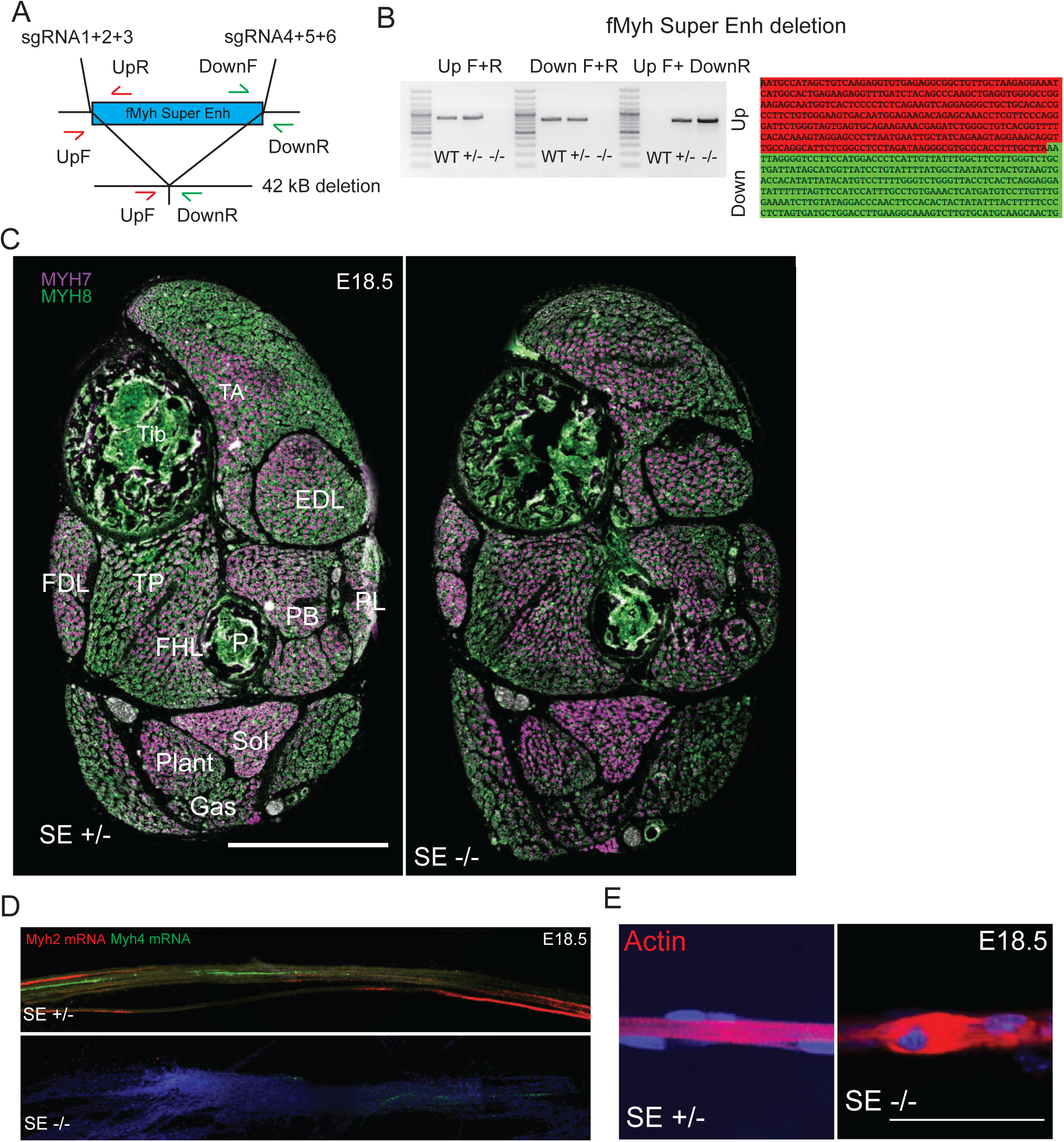
Phenotype of f*Myh*-SE deletion in E18.5 mutant fetuses. (A) Diagram showing the strategy for *in vivo* deletion of the f*Myh-SE* by CRISPR/Cas9. (B) Sequence of the deleted allele. (C) Immunostaining of distal hindlimb sections of control and *fMyh-SE^-/-^* showing MYH8 (green), MYH7 (purple) and Laminin (white) expression. (D) RNascope experiments against *Myh2* and *Myh4* mRNAs on isolated E18.5 fibers of control and mutant fetuses. (E) Myofibers from mutant diaphragm showed defects in sarcomeres organization as revealed by phalloidine staining. For E, scale bar: 500 μm.

**Figure S8.**
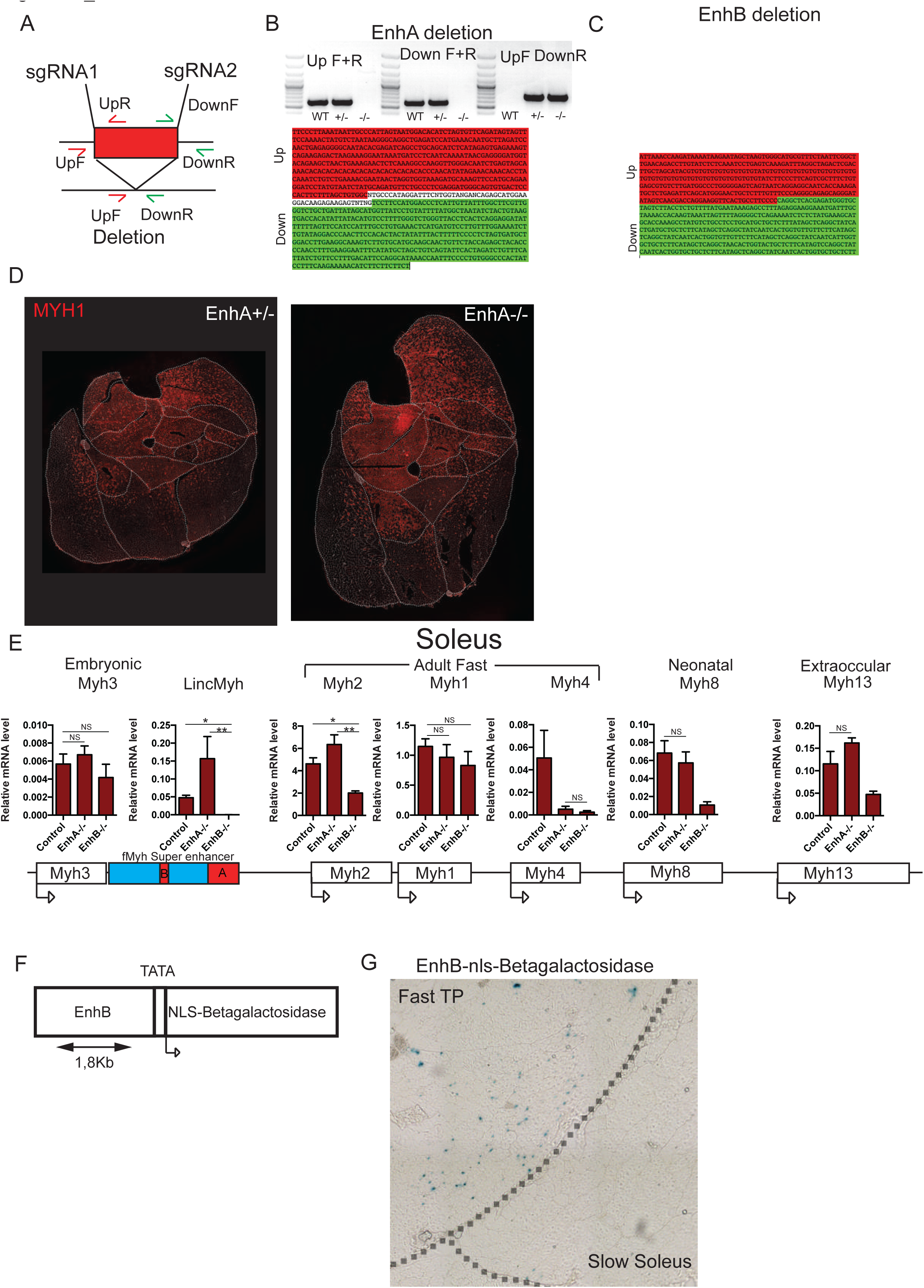
Muscle phenotype of enhancers A and B deletion. (A) Diagram showing the strategies for *in vivo* deletions of enhancer A or B by CRISPR/Cas9. (B) PCR showing the DNA fragment amplified after *EnhA* deletion, and sequence of the deleted allele. (C) Sequence of the deleted *EnhB* allele. (D) Immunostaining against MYH1 on adult distal hindlimb sections in control and *EnhA^-/-^*. (E) Quantification of *Myh3, Myh2, Myh1, Myh4, Myh8* and *Myh13* mRNAs and of *Linc-Myh* by RT-qPCR in Soleus of control, *EnhA* and *EnhB* mutants. (F) Schema of the transgene used to generate a transgenic mouse line expressing nuclear beta-galactosidase under the control of the 1.8kb *EnhB* DNA element. (G) Beta-galactosidase positive nuclei are detected in the fast tibialis posterior but not in the slow soleus. For E, n=3. Numerical data are presented as mean ± S.E.M. \**P* < 0.05, \*\**P* < 0.01, \*\*\**P* < 0.001

**Figure S9.**
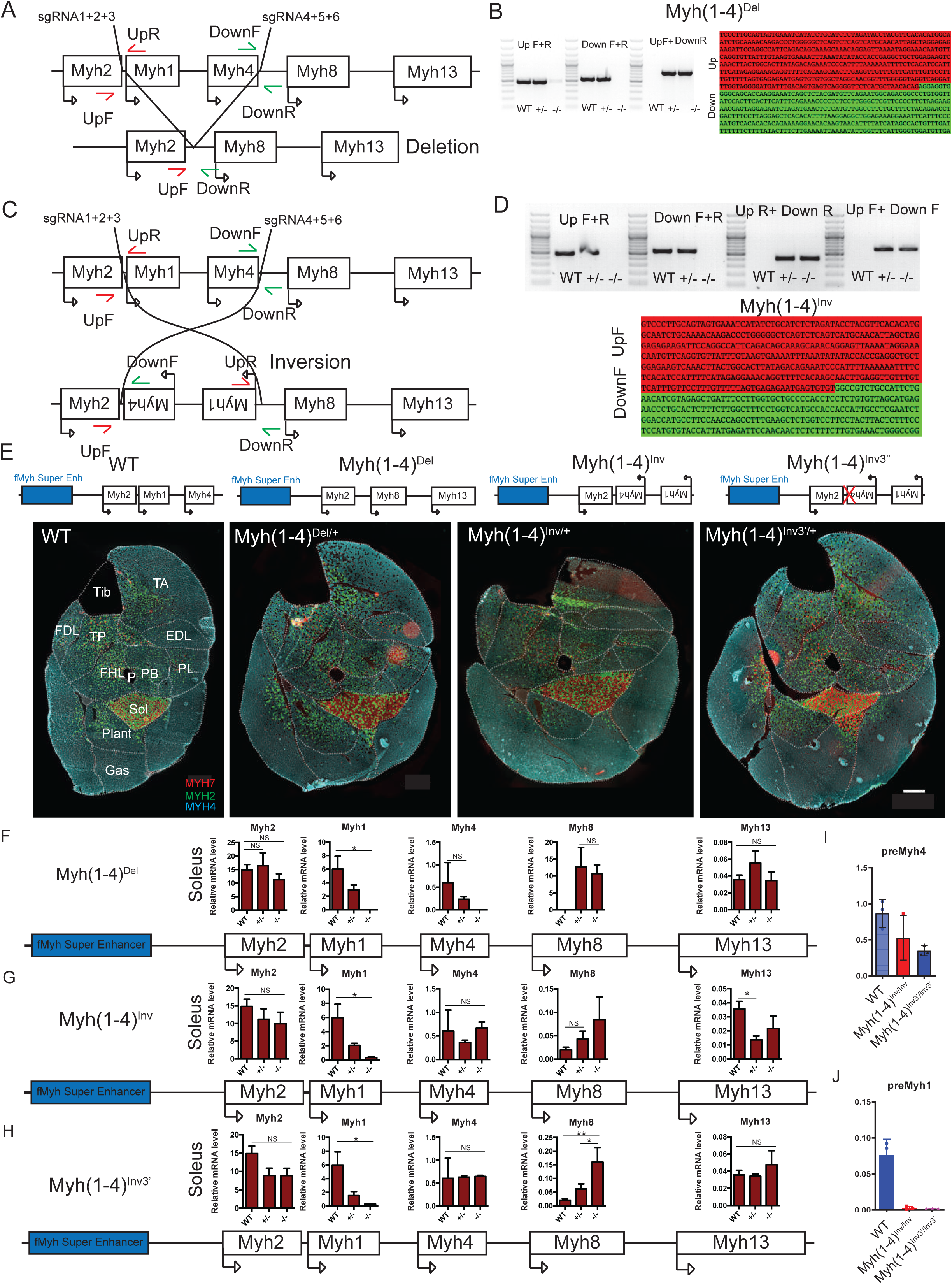
Competition of the f*Myh* gene promoters for the shared SE. (A) Diagram showing the strategy for *in vivo* deletion of *Myh1* and *Myh4* by CRISPR/Cas9. (B) PCR used for the screening of mutant animals and sequence of the deleted allele. (C) Diagram showing the strategy for *in vivo* inversion of *Myh1* and *Myh4* by CRISPR/Cas9. (D) PCR used for the screening of mutant animals and sequence of the inverted allele. (E) Immunostaining against MYH2, MYH4 and MYH7 proteins of adult distal hindlimb sections of 2-3-month-old WT, *Myh(1-4)^Del/+^*, *Myh(1-4)^Inv/+,^* and of *Myh(1-4)^Inv3’/+^* animals. (F) Quantification of *Myh2, Myh1, Myh4, Myh8* and *Myh13* mRNAs by RT-qPCR from adult Soleus of WT, *Myh(1-4)^Del/+^*, and *Myh(1-4)^Del/Del^* animals. (G) Quantification of *Myh2, Myh1, Myh4, Myh8* and *Myh13* mRNAs by RT-qPCR from adult Soleus of WT, *Myh(1-4)^Inv/+^* and *Myh(1-4)^Inv/Inv^* animals. (H) Quantification of *Myh2, Myh1, Myh4, Myh8* and *Myh13* mRNAs by RT-qPCR from adult Soleus of WT, *Myh(1-4)^Inv3’/+^* and of *Myh(1-4)^Inv3’/Inv3’^* animals. (I) Quantification of *Myh4* premRNA from adult TA of WT, *Myh(1-4)^Inv/Inv^* and of *Myh(1-4)^Inv3’/Inv3’^* animals. (J) Quantification of *Myh1* premRNA from adult TA of WT, *Myh(1-4)^Inv/Inv^* and of *Myh(1- 4)^Inv3’/Inv3’^* animals. Numerical data are presented as mean ± S.E.M. \**P* < 0.05, \*\**P* < 0.01, \*\*\**P* < 0.001

**Table S1.** Antibodies used in the study.

| Target | Antibody reference | Supplier | Species | dilution |
| --- | --- | --- | --- | --- |
| MYH7 | BA-F8 | DHSB | mouse IgG2B | 1/40 |
| MYH2 | SC-71 | DHSB | mouse IgG1 | 1/200 |
| MYH1 | 6H1 | DHSB | mouse IgM | 1/40 |
| MYH4 | BF-F3 | DHSB | mouse IgM | 1/200 |
| MYH8 | N3.36 | DHSB | mouse IgM | 1/200 |
| MYH13 | 4A6 | DHSB | mouse IgM | 1/200 |
| MYH3 | Bf-45 | DHSB | mouse IgG1 | 1/200 |
| Laminin | L9393 | Sigma | Rat | 1/500 |
| GFP | ab290 | Abcam | Chicken | 1/200 |
| Neurofilament | 841001 | BioLegend | Rabbit | 1/1000 |
| alpha bungarotoxin | B13422 | Thermofischer |  | 1/500 |

**Table S2.**
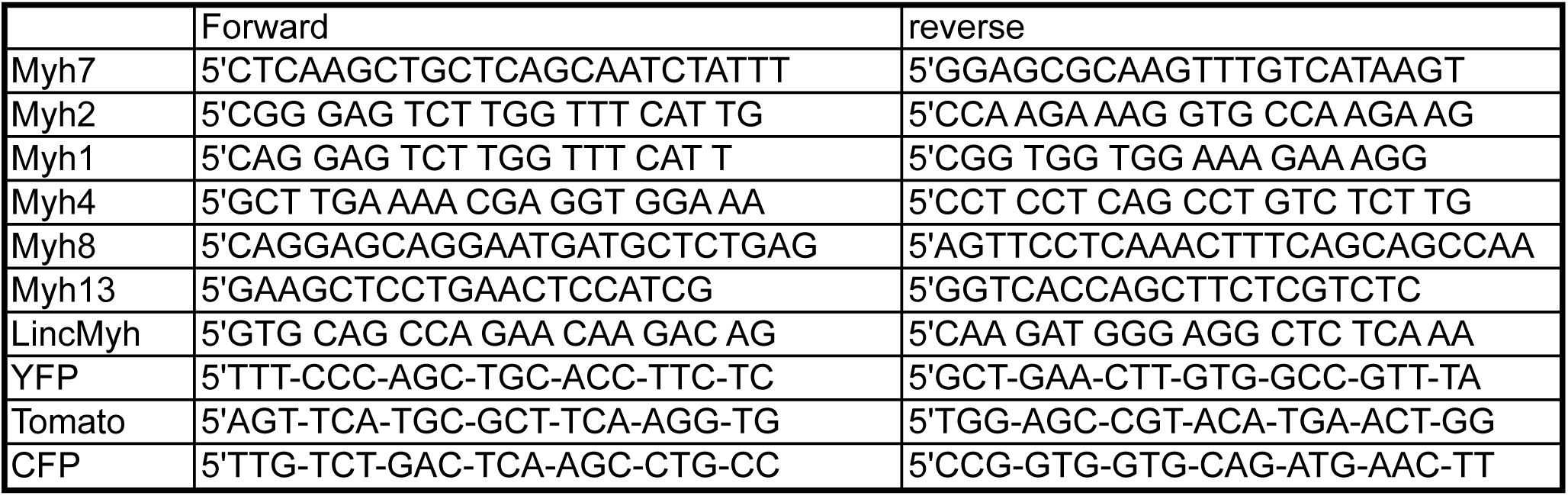
Oligonucleotides used for RT-qPCR experiments.

**Table S3.**
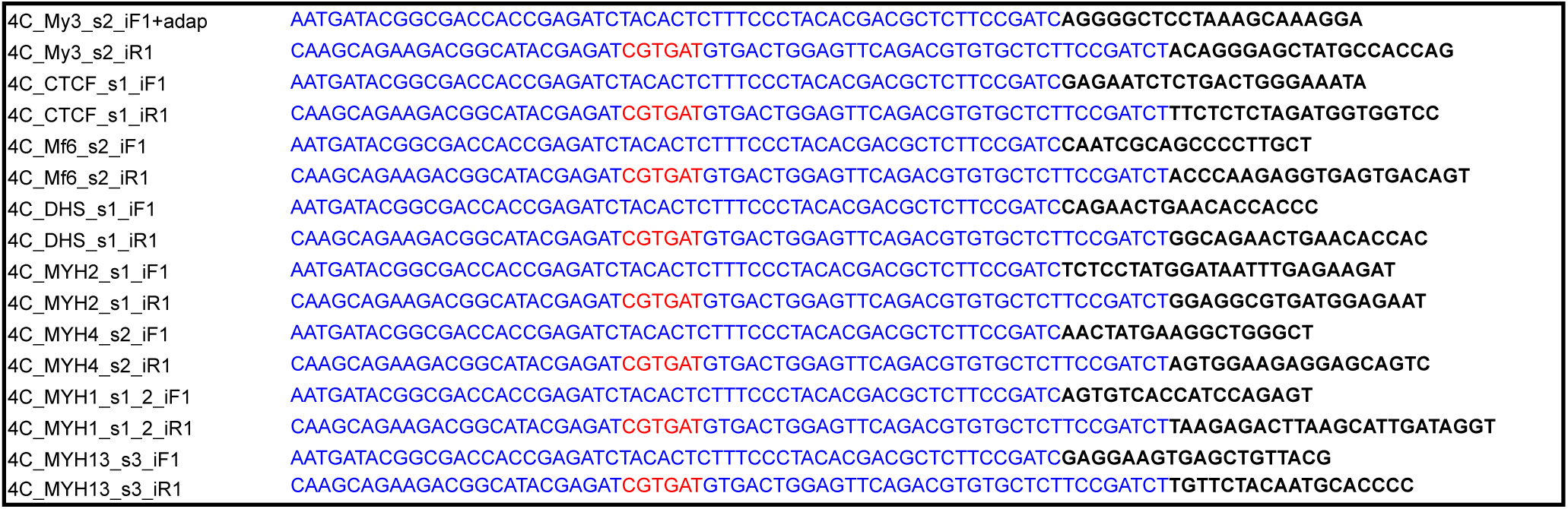
Oligonucleotides used for 4C-seq experiments.

**Table S4.**
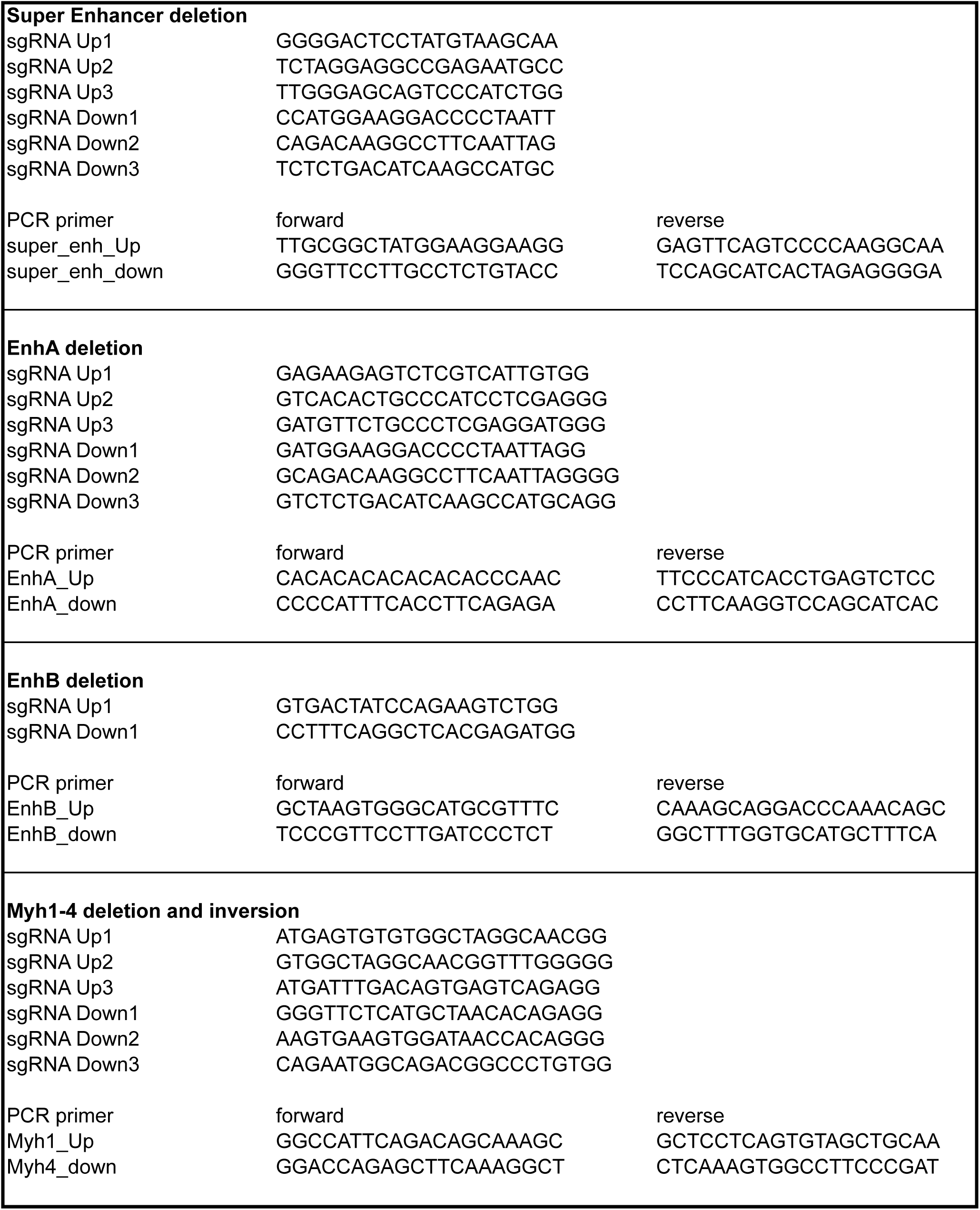
sgRNA and PCR primers used for screening of CRISPR/Cas9 edited genome.

